# Quiescence improves *Candida albicans* survival of fungicidal drug exposure

**DOI:** 10.64898/2026.02.27.708532

**Authors:** Ozan Berk Imir, Marina Druseikis, Ying Xie, Yujia Ji, Liam Holt, Judith Berman, David Gresham

**Author notes:** equal contribution (Ozan Berk Imir, Marina Druseikis).

## Abstract

Quiescence, defined as the reversible exit from mitotic division and proliferative growth, is the predominant state of all microbes. Despite its prevalence, the properties and consequences of quiescence in *Candida albicans*, an opportunistic fungal pathogen, remain largely unexplored. In this study, we characterized the morphological, molecular, and biophysical properties of quiescent *C. albicans* cells and assessed the effects of quiescence on antifungal drug efficacy. Quiescent cells that were induced via carbon starvation in rich and minimal media underwent distinct morphological changes upon entry into quiescence; this included an increase in cell buoyant density, altered fluidity of the cytoplasm and nucleus, and remodeling of mitochondria. Most *C. albicans* cells arrested in an unbudded G1/G0 state, although a significant fraction of cells had budded morphologies and 4N DNA content, indicating that they arrested at other cell cycle phases. Both budded and unbudded quiescent cells efficiently re-entered the cell cycle upon nutrient replenishment, with time-to-quiescence exit varying depending on the total nutritional quality of the medium. Quiescence was associated with large-scale gene expression remodeling, including downregulation of ribosomal biogenesis genes and upregulation of autophagy and stress response pathways. Notably, a greater proportion of quiescent cells than proliferative cells survived exposure to the commonly used antifungal drugs micafungin, caspofungin, and amphotericin B in genetically diverse strains. Thus, quiescence is a distinct cellular state with important implications for antifungal drug efficacy in *C. albicans*.

**Author Summary:** We show that *Candida albicans*, a common fungal pathogen, can enter a reversible, non-dividing state when starved of carbon. Starved cells become smaller and denser, reorganize their mitochondria, change how densely packed the inside of the cell and its nucleus are, and switch on stress-protection and internal recycling programs while reducing protein synthesis activity. Most cells have ceased to actively divide, but many retained budded shapes and could restart growth when nutrients returned; the timing of recovery depended on the nutritional environment in which quiescence was initiated. Critically, quiescent cells from laboratory and clinical strains exhibited greater survival than proliferative cells when exposed to widely used fungicidal drugs including micafungin, caspofungin, and amphotericin B. These findings indicate that quiescence is an active, adaptive physiological state that helps *Candida albicans* survive hostile environmental conditions such as temperature stress and drug exposure. Accounting for the metabolic state of fungal cells in diagnostics and drug development may improve treatment outcomes.

## Introduction

*Candida albicans* is one of the most common human fungal pathogens. Although typically a harmless commensal resident of the oral, gastrointestinal, and genital flora (1,2), *C. albicans* is an opportunistic pathogen (3–5) that causes infections ranging from superficial mucosal infections to life-threatening invasive candidiasis (6,7). Worldwide, *C. albicans* causes approximately 150 million infections and over 100,000 deaths annually, and thus, the World Health Organization classified *Candida albicans* and a related fungus, *Candida auris,* as critical priority fungal pathogens due to the increasing occurrence of drug resistance and treatment challenges (8,9).

Effective treatment of microbial infections requires an understanding of their molecular and physiological properties. In response to nutrient deprivation signals, many microbes can reversibly exit the cell cycle and enter quiescence, a non-proliferative state that enhances survival in unfavorable conditions (10,11). In bacteria, quiescence is synonymous with dormancy, a survival strategy employed to evade immune responses and minimize the metabolism of antimicrobial drugs (12,13). In model eukaryotic microbes, such as *Saccharomyces cerevisiae* and *Schizosaccharomyces pombe*, quiescence involves extensive reprogramming, including activation of regulated gene expression programs, increased autophagy, and chromatin reorganization that enables long-term viability in the absence of active proliferation (14–16).

In eukaryotic microbes, quiescence is regulated by evolutionarily conserved signaling pathways including the Target of Rapamycin (TOR) and AMP-activated protein kinase (AMPK), which coordinate cellular responses to environmental signals (17,18). The state of quiescence in these eukaryotes is associated with increased resistance to high temperature, oxidative stress, and DNA damage (18–20), suggesting that increased stress resistance is a general characteristic of quiescent cells. Whether *C. albicans* can establish quiescence in response to nutrient starvation signals, and the consequences of quiescence for stress resistance, have not been investigated previously.

In addition to increased resistance to common environmental stresses, the lower metabolic activity of quiescent cells may have important consequences for therapeutic strategies using antifungals. Echinocandins, such as caspofungin and micafungin inhibit the synthesis of β-1,3-D-glucan, an essential component of the fungal cell wall (21), whereas polyenes, such as amphotericin B bind membrane ergosterol to form pores that drive ion leakage and cell death (22). Disruption of these structural components compromises cell integrity, leading to osmotic instability, lysis, and cell death (23). As many metabolic and synthetic cellular processes are down-regulated in quiescence, we hypothesized that cytotoxic compounds might be less effective in killing quiescent cells.

Here, we found that *C. albicans* effectively initiates a quiescent state in response to carbon starvation, undergoing physiological and morphological remodeling at the cellular level and large-scale changes in gene expression and biophysical properties at the molecular level. In both rich and minimal media, quiescent cells were more dense, underwent mitochondrial remodeling, and were more resistant to heat stress. Using flow cytometry, rheological tracking, and single-cell microscopy, we find that the majority of starved cells grown in rich media arrest growth in G1 of the cell cycle. However, there was significant inter-individual heterogeneity in cell cycle arrest morphology, especially in cells starved in minimal media, indicating that arrest in G1 is not a requirement for quiescence. Interestingly, carbon-starved cells exhibited reduced cytoplasmic and nuclear fluidity when starved in rich medium, but increased cytoplasmic and nuclear fluidity when starved in minimal media. Transcriptomic profiling indicates that quiescent cells undergo broad metabolic reprogramming characterized by downregulation of biosynthetic pathways and upregulation of stress response genes. Finally, quiescent cells survived exposure to fungicidal drugs better than proliferative cells. Thus, the quiescent cell state challenges effective antifungal drug efficacy, motivating further investigation into the regulation of quiescence in fungal pathogens and strategies for mitigating its influence on drug responses.

## Results

### Physiological properties of quiescent *C. albicans*

Microbial population growth is characterized by a lag phase, an exponential growth phase, and a stationary phase in which population growth ceases (**Figure 1A**) due to the depletion of one or more essential nutrients. Identification of the nutrient that limits growth capacity is essential, as starvation of different nutrients can elicit distinct cellular processes and require different signaling pathways. Therefore, we asked if carbon is the limiting nutrient in (i) rich media (YPD) and (ii) defined synthetic minimal media, using a two-fold dilution series of glucose concentrations with all other media components unchanged (**Figure 1B**). The relationship between carbon concentration and final culture yield was linear for both rich (**Figure 1C**) and minimal (**Figure 1D**) media, consistent with carbon starvation being the initiating signal for growth arrest in these conditions. For all subsequent experiments, we used 2% glucose (0.11 M carbon) in rich media and 0.05% glucose (0.00167 M carbon) in minimal media. Cells growing exponentially and in stationary phase were morphologically similar, although cell size was significantly smaller in rich media stationary phase cultures (**Figure 1E**). As cultures may be heterogeneous, we defined “proliferative-enriched” samples as cultures sampled at 6 hours post-inoculation (“proliferative” hereafter) and “quiescent-enriched” samples as cultures sampled at 72 hours post-inoculation (“quiescent” hereafter).

**Figure 1.**
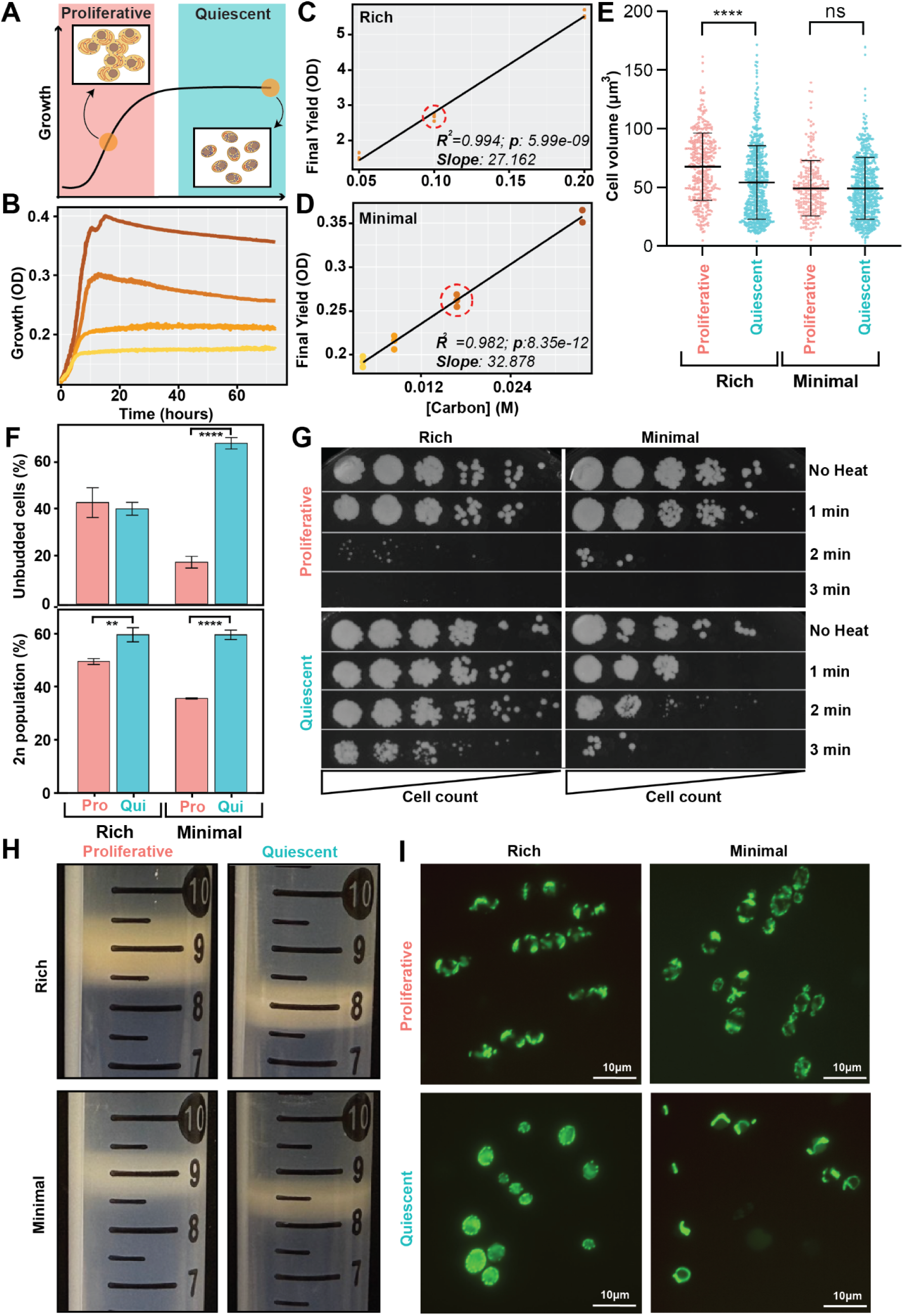
Properties of quiescent *Candida albicans*. **A)** Definition of microbial growth phases. **B)** Population growth profiles of *Candida albicans* quantified using absorbance (OD at 600nm) in minimal media with variable initial concentrations of glucose: 5.6 mM (n = 2), 2.8 mM (n = 3), 1.4 mM (n = 3), and 7 mM (n = 3). **C)** Final population yield (OD at 600nm) versus initial glucose concentration for rich media. The line indicates a linear model fit to the data. The dashed circle indicates the concentration of glucose used in rich media for all subsequent experiments. **D)** Final population yield (OD at 600nm) versus initial glucose concentration in minimal media. The line indicates a linear model fit to the data. The dashed circle indicates the concentration of glucose used in minimal media for all subsequent experiments. **E)** Quantification of cell volume for cells grown in rich and minimal media for proliferative and quiescent cells.The median value is reported in each panel and indicated by the horizontal line ± interquartile range. The p-value was determined using Student’s t-test, ∗∗∗∗p < 0.0001, ns: p > 0.05 **F)** Bud index quantification (top) and DNA content staining tusing SYTOX Green and flow cytometry(bottom). **G)** Cells were subjected to 1, 2, or 3 minutes of 56 °C temperature stress. 5-fold serial dilutions of cells were transferred using a pin frogger and incubated for 2 days prior to imaging. **H)** Percoll-based density fractionation of proliferative and quiescent cells. Denser cells migrate further through the gradient and thus appear as lower bands. **I)** Mitochondrial visualization using a mNeonGreen-SFC1 protein fusion. Cells were excited with a 470 nm wavelength; scale bar indicates 10 µm.

In *S. cerevisiae,* nutrient starvation typically results in cell cycle arrest in the G1/G0 stage of the cell cycle (11), although *S. cerevisiae* and *S. pombe* can reversibly initiate quiescence at any point of the cell cycle (24,25). For *C. albicans,* the cell cycle stage was quantified both as the fraction of unbudded cells and as the DNA content. Initially, unbudded cells made up 30-40% of proliferative cells in both rich and minimal media, increasing to 60-70% in quiescent cells grown in minimal media. Surprisingly, cells grown in rich media had exhibited a more heterogeneous budding profile when quiescent (**Figure 1F** and **Supplementary Figure 1**). DNA content staining largely recapitulated the morphological analysis: in growing cultures, approximately 60-70% of cells had a 4N (G2/M phase) DNA content; this decreased with time (**Supplementary Figure 2**) to 35-40% of starved cells (**Figure 1F**). These assays suggest that 60-70% of quiescent *C albicans* cells are arrested in the G1/G0 phase of the cell cycle when starved for carbon.

To ask if quiescence confers increased thermotolerance in *C. albicans,* we evaluated heat killing in proliferative and quiescent cells by plating serial dilutions of cells subjected to 56°C heat shock for increasing periods of time. In both rich and minimal media, quiescent cells survived longer periods of high heat exposure than proliferative cells (**Figure 1G**), indicating that quiescence confers increased heat stress tolerance.

*S. cerevisiae* cells become denser over time when they initiate quiescence. The density of proliferative and quiescent *C. albicans* cells, therefore, was determined by density fractionation with Percoll gradients. In both rich and minimal media, quiescent cells were more dense than proliferative cells (**Figure 1H** and **Supplementary Figure 3**). The difference in density between proliferative and quiescent cells was larger for rich medium than for minimal medium, which may result from the decrease in cell size in rich media. Thus, quiescent cells in both rich and minimal media are denser than proliferative cells.

#### Mitochondrial structure differs between proliferative and quiescent cells

In *S. cerevisiae,* mitochondrial reorganization is a hallmark of quiescence (14). We first tried to assess mitochondrial morphology in live quiescent *C. albicans* cells using Mitotracker Red FM. In proliferative cells, the expected tubular mitochondrial network spanned from the mother to the bud (**Supplementary Figure 4**) in both rich and minimal media. By contrast, Mitotracker Red FM stained a vacuole-like structure in cells grown in rich and minimal medium were either unstained or became very bright, a signal usually associated with dead cells (**Supplementary Figure 4**). Similar staining patterns were seen with two different clinical isolates, suggesting that this is a feature of quiescent cells rather than strain background (**Supplementary Figure 5)**. We hypothesize that quiescent cells have altered membrane permeability to Mitotracker Red FM stain making it an inappropriate assay in quiescent cells.

We reasoned that using a strain expressing a fluorescently labeled protein that localizes to mitochondria would circumvent the inability to visualize mitochondria in quiescent cells. We expressed the gene encoding a mitochondrial succinate-fumarate transporter fusion protein (*mNeonGreen-SFC1*) in *C. albicans* cells and starved them for 72 hours prior to visualization by fluorescence microscopy. As expected, proliferative cells grown in rich or minimal media that expressed the mNeonGreen-SFC1 fusion protein co-localized with Mitotracker Red FM to a tubular mitochondrial network spanning from mother to bud (**Supplementary Figure 6**) in both minimal and rich media (**Figure 1I**). By contrast, the majority of quiescent cells starved in rich media exhibited a vesicular or globular mitochondrial network (yellow arrows) and an overall increase in the mNeonGreen-SFC1 fusion protein signal. In addition, some of the quiescent cells starved in rich medium had fragmented tubular morphologies (cyan arrows). In minimal medium, the morphological changes in quiescent cells included a tubular morphology that was less pronounced than in proliferative cells (**Figure 1l**) and some cells that did not show any mitochondrial structures and a highly reduced or completely absent mNeonGreen-SFC1 fusion protein signal, possibly indicating senescence or death (white arrows) (**Figure 1l**) (**Supplementary Figure 7**). Taken together, these results suggest that quiescent *C. albicans* cells undergo mitochondrial reorganization, and that the extent of mitochondrial reorganization is more dramatic in rich media than in minimal media.

### Gene expression profiling

To quantify gene expression changes during the transition from proliferating to quiescent cultures, we performed RNA sequencing and comparative transcriptomic analysis in rich and minimal media at multiple timepoints (**Figure 2A**). Functional annotation of hierarchically clustered gene expression changes over time indicated both metabolic and regulatory remodeling of the transcriptional program (**Figure 2B**). We defined 7 distinct expression clusters (cluster 1-7) with largely consistent trends in both media, and one cluster (cluster 8) that had divergent expression patterns in the two media. Significant over-representation of Gene Ontology (GO) terms was used to annotate functions associated with the clusters (**Supplementary Figure 8** and **Figure 2B**).

**Figure 2.**
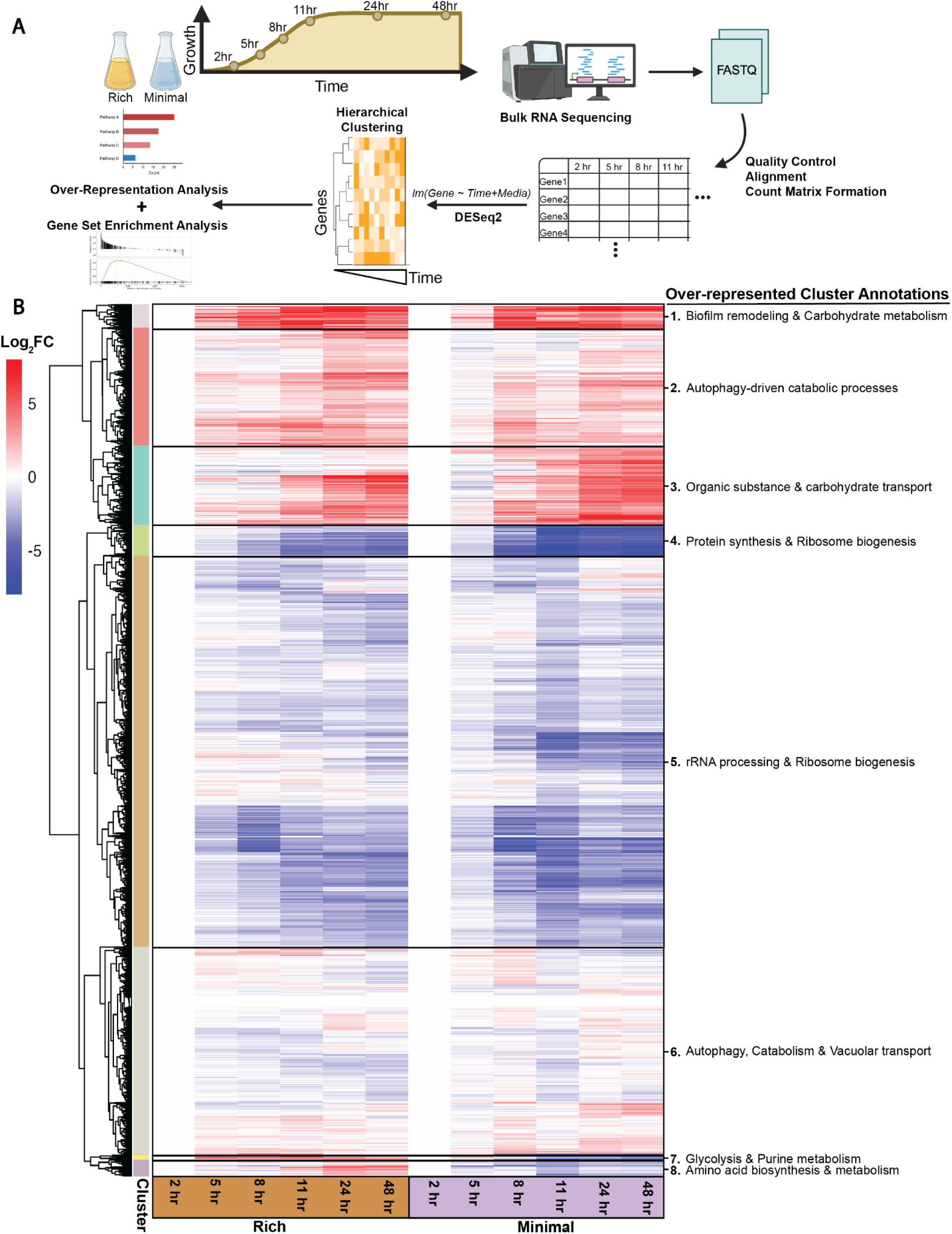
Gene expression profiling in proliferative and quiescent *Candida albicans* cells. **A)** Experimental design, sequencing pipeline, and computational strategy employed to quantify transcript level changes across time. **B)** Hierarchical clustering defined 8 clusters that are annotated using the most significant over-represented GO terms (p < 0.05) (**Supplementary Figure 7**).

Genes that increased in expression with the establishment of quiescence were enriched for biofilm formation and carbohydrate metabolism (cluster 1), autophagy-driven catabolism (cluster 2), organic-substance and carbohydrate transport (cluster 3), and autophagy/catabolism with vacuolar transport (cluster 6). Conversely, genes involved in protein synthesis and ribosome biogenesis were repressed (clusters 4–5), as were pathways for glycolysis and purine metabolism (cluster 7). These results are generally consistent with a broad cessation of growth-associated processes and metabolic remodeling in quiescent cells. Cluster 8, which was enriched for amino-acid biosynthesis and metabolism pathways, was unique in that the expression of genes in these pathways increased in rich medium and decreased in minimal medium over time. This likely reflects the very different level of non-carbon nutrient availability in the two media. Furthermore, translation-related processes tended to be more highly expressed in rich media at all time points (**Supplementary Figure 9**). This may reflect the increased protein production capacity associated with nutrient-rich environments that persists even as starvation commences and quiescence is established.

### Cytoplasmic and nuclear fluidity

We next asked if there were differences in the biophysical intracellular properties of proliferating and quiescent *C. albicans* cells. We used <u>g</u>enetically <u>e</u>ncoded <u>m</u>ultimeric nanoparticles (GEMS) with a diameter of 40nm (40nm-GEMs) localized to either the cytosol (**Figure 3A**) or the nucleus (**Figure 3B**). GEMs are brightly fluorescent mesoscale particles, which allows measurement of the crowding of the cytoplasm or nucleus, expressed as the effective diffusion coefficient (*D*_eff_) (26). Cells were imaged every 10 milliseconds for 2 seconds to obtain trajectories for single particle tracking. We observed comparable continuous trajectory lengths in both proliferating and quiescent cells, with median values of 22 frames in the cytosol (**Supplementary Figure 10A**) and 18 frames in the nucleus (**Supplementary Figure 10B**). The effective diffusion coefficient (*D*_eff100ms_) of 40nm-GEMs was calculated based on mean-square displacement over the first 10 timeframes, corresponding to the first 100ms of acquisition (**Supplementary Figure 10C**). At least three trajectories per cell were analyzed.

**Figure 3.**
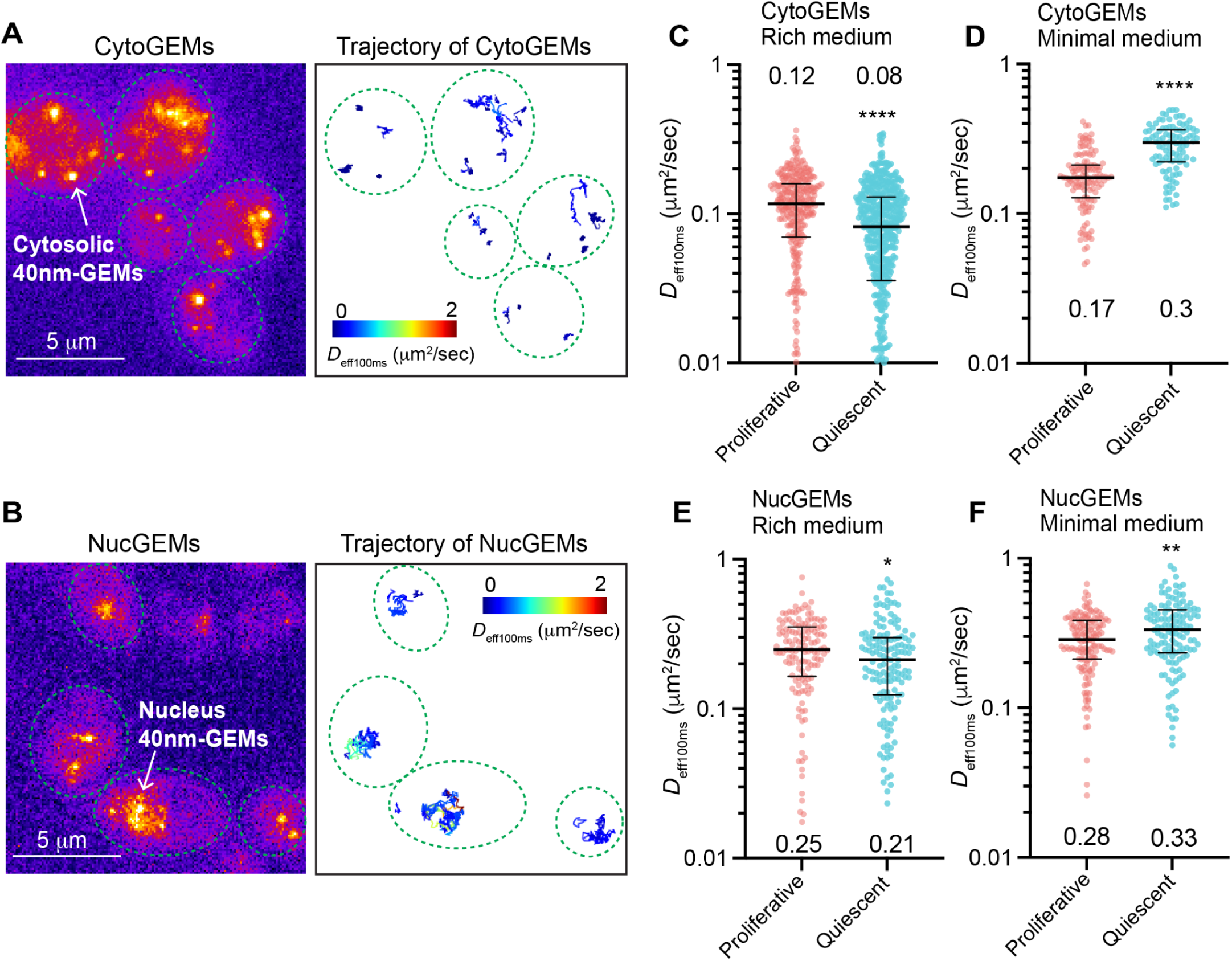
Intracellular fluidity differs depending on media composition, growth state and subcellular compartment in *C. albicans*. Representative live cell image of **A)** cytosolic and **B)** nuclear 40nm-GEMs expressed in *Candida albicans*. Single particle tracking of 40nm-GEMs was performed and trajectories determined for individual GEMs to quantify effective diffusion rates. Multiple GEMs (>3) were tracked within each cell. **C)** The effective diffusivity at 100 ms (D_eff100ms_) for 40nm-GEMs localized to the cytoplasm in proliferating and quiescent *C. albicans* cells cultured in rich media. Proliferative: n = 324 cells; quiescent: n = 515 cells. **D)** The effective diffusivity at 100 ms for 40nm-GEMs localized to the cytoplasm in proliferating and quiescent *C. albicans* cells grown in minimal media. Proliferative: n = 127 cells. quiescent: n = 104 cells. **E)** The effective diffusivity of 40nm nuclear GEMs in proliferating and quiescent *C. albicans* cells cultured in rich media. Proliferative: n = 139 cells; quiescent: n = 149 cells. **F)** The effective diffusivity of 40nm nuclear GEMs in proliferating and quiescent *C. albicans* cells cultured in minimal media. Proliferative: n = 146 cells; quiescent: n = 134 cells. Each point represents an individual cell. The median value is reported in each panel and indicated by the horizontal line ± interquartile range. The p-value was determined using Student’s t-test, ∗p < 0.05, ∗∗p < 0.01, ∗∗∗p < 0.001, ∗∗∗∗p < 0.0001).

In rich media, quiescent cells had less cytoplasmic fluidity than proliferating cells (**Figure 3C**). Surprisingly, in minimal media, quiescent cells had more cytoplasmic fluidity than proliferating cells (**Figure 3D**). This distinction between media conditions was also evident when we used 40nm-GEMs that localize to the nucleus: nuclear 40nm-GEMs *D*_eff100ms_ decreased slightly but significantly (p<0.05) in quiescent cells starved in rich media (**Figure 3E**). By contrast, nuclear 40nm-GEMs *D*_eff100ms_ increased significantly (p<0.01) in quiescent cells starved in minimal media (**Figure 3F**). Thus, in quiescent cells, both cytoplasmic and nuclear fluidity decreased in rich media but increased in minimal medium.

Macromolecular crowding levels are influenced by macromolecule quantity, size, and intracellular free space. Since cell volume is a strong determinant of intracellular free space, we measured cell volume and examined its correlation with fluidity in individual cells. A weak positive correlation between nuclear fluidity and cell volume was measured in minimal media for both proliferating and quiescent cells; this was not apparent in rich media, suggesting that cell volume cannot fully explain the difference in altered fluidity that occurred in the different culture media (**Supplementary Figure 11)**.

Culture media pH has a strong impact on intracellular fluidity and dormancy in *S. cerevisiae* and *S. pombe* (27) when energy is depleted. Therefore, we asked if culture media pH potentially contributed to the divergent quiescent fluidity responses of *C. albicans* cells. Interestingly, after 72 h of culturing, the average pH of rich media increased from pH 5.5 to pH 7.2, whereas the average pH of minimal media decreased from pH 4.5 to pH 3.0 (**Supplementary Figure 12**). Thus, media pH may strongly affect the physiology of quiescent *C. albicans* cells. The apparent reduced diffusivity in proliferative cells grown in rich media compared with minimal media may be a result of differences in pH or alternatively, could reflect the increased translational capacity of cells grown in rich media as ribosomes are a key contributor to molecular crowding in the cytoplasm (26).

### The dynamics of exit from quiescence depend on environmental conditions

The defining feature of quiescent cells is their capacity to reinitiate cell growth and division once nutrient limitation is relieved. Experimentally, this is accomplished by providing cells with fresh medium. To study exit from quiescence we starved cells in either rich or minimal media and then replenished them with rich media (YPD 2% glucose). We used a microcolony growth assay (28) to monitor the return-to-growth of cells sampled from both proliferative (log phase) and quiescent (72 h) cultures. We sought to determine whether those cells that had arrested as unbudded cells or with one or more buds were able to reinitiate growth and division upon refeeding.

All cells, including unbudded, single budded, and multi-budded cells, were monitored over 9 hours of refeeding to determine the proportion, and timing, of cells that could initiate cell division **(Figure 4A)**. The majority of cells isolated 72 hours after inoculation reinitiated cell division (95% from rich media and 94% from minimal media) and thus, by definition, were quiescent (**Figure 4B** and **Figure 4C**). From proliferative cultures, 97-100% of the cells initiated budding within 1 hour of refeeding (**Figure 4B** and **Figure 4C**). In rich medium, the majority of quiescent cells were unbudded (65%); in minimal medium the majority of cells (74%) had at least one bud prior to refeeding (**Figure 4D**), and the number of budded cells appeared to decrease with time in culture prior to refeeding **(Figure 1F**, **Figure 4D** and **Figure 4E)**. To test the generalizability of these observations, we asked about the degree of unbudded cells and the timing of return to growth for an evolutionarily distant clinical isolate, P34048. Importantly, the degree of budding was similar for P34048 and SC5314 in both rich and minimal media (**Figure 4E**). Thus, the ability of cells to arrest as budded or unbudded cells during starvation is not exclusive to the lab strain.

**Figure 4.**
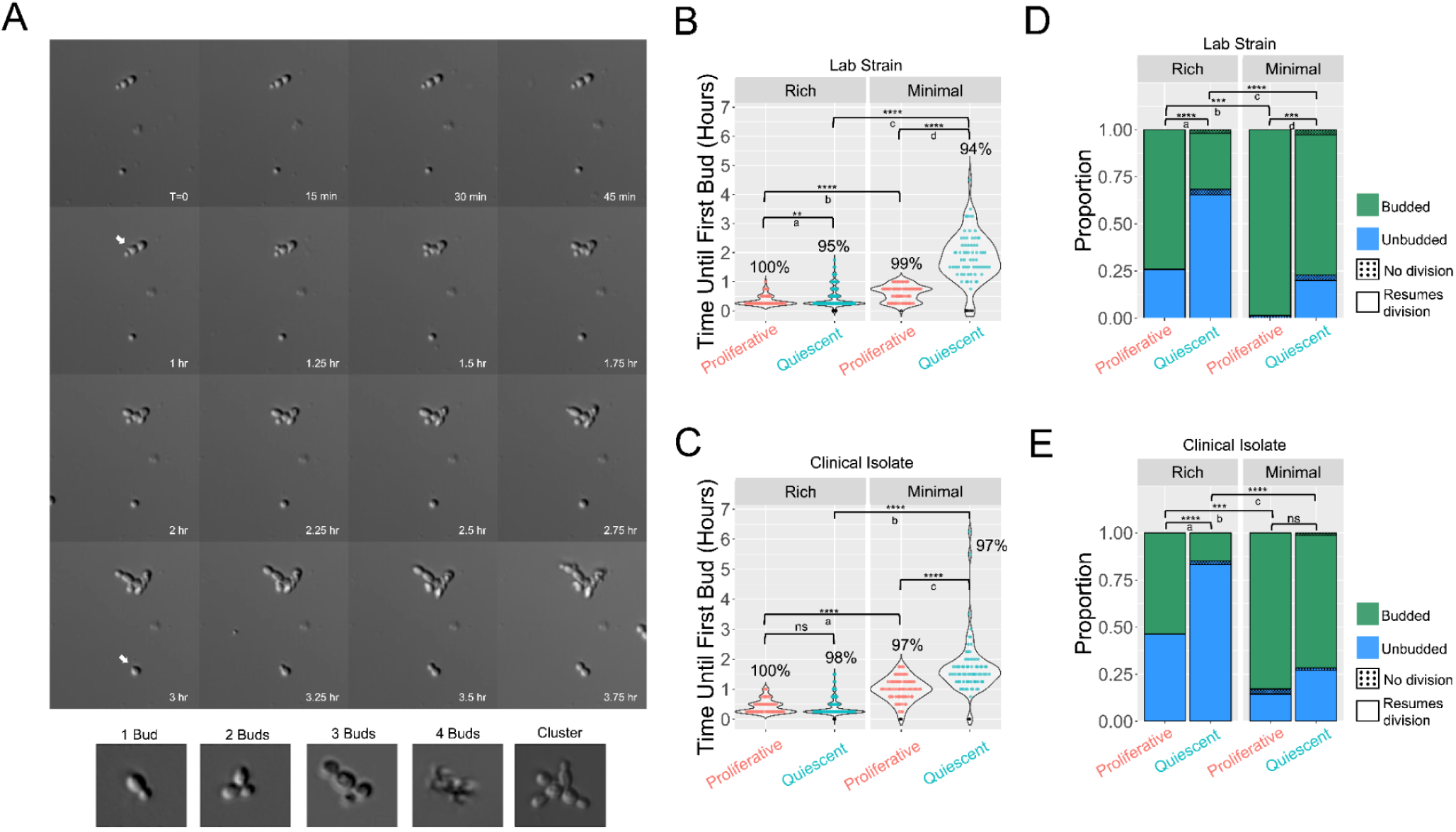
Quiescent cells reinitiate cell growth and division. We quantified cell budding dynamics using a microcolony growth rate assay in proliferative and quiescent culture. **A)** Representative time-lapse microscopy images from the microcolony assay and examples of different types of budding states in starved cultures (not to scale). White arrows denote examples of bud emergence. **B)** Time required until first bud emergence of cells from proliferative and quiescent cultures grown in rich media and minimal media in the lab strain SC5314. We defined the time until first bud as the time required for a new bud to be detected on a cell. A score of 0 indicates that a cell did not produce a bud within the 9-hour timeframe (indicated as a black dot). The percentage of cells that resumed cell division is denoted above each violin plot. p-values for: a) 0.00191; b) 8.30E-10; c) 6.51E-22; d) 3.27E-19. **C)** Time required until first bud emergence of cells from proliferative and quiescent cultures grown in rich media and minimal media in the clinical isolate P34048. p-values for: a) 5.67E-19; b) 6.80E-20; c) 1.19E-08. **D)** Budding status and resumption of division of in lab strain SC5314. Dotted bars indicate the proportion within a specific budding status that did not produce a new bud within the 9-hour time frame. p-values for a) 7.32E-09; b) 0.000105; c) 2.83E-11; d) 0.00033. **E)** Budding status and resumption of division of P34048. Dotted bars indicate the proportion within a specific budding status that did not produce a new bud within the 9-hour time frame. p-values for a) 7.32E-09; b) 0.000649; c) 3.95E-19. SC5314 rich proliferative, n = 58; SC5314 rich quiescent, n = 243; SC5314 minimal proliferative, n = 75; SC5314 minimal quiescent, n = 70; P34048 rich proliferative, n = 54; P34048 rich quiescent, n = 208; P34048 minimal proliferative, n = 76; P34048 minimal quiescent, n = 74. p-values for significant differences between conditions or cultures are denoted with asterisks. For **B** and **C**, The p-values were determined using Student’s t-test; for **D** and **E**, the p-values were determined using Chi-squared test with Benjamini–Hochberg correction. ∗p < 0.05, ∗∗p < 0.01, ∗∗∗p < 0.001, ∗∗∗∗p < 0.0001).

We measured the time at which cells in a starved culture initiated division (detected as the first appearance of a new bud (**Figure 4A**). Interestingly, the dynamics of reinitiation of growth (i.e. exit from the quiescent state when provided with rich medium) differed by ∼2.5-fold between cells starved for 72 hours in rich versus minimal media: the time required for all quiescent cells to bud was within 1.75 versus 4.5 hours, respectively **(Figure 4B)**. Thus, the dynamics of cell division following starvation depends on the composition of the media in which quiescence was initiated. Furthermore, the time required for budding was similar for clinical strain P34048 as for lab strain SC5314 in both rich and in minimal media (**Figure 4B** and **Figure 4C**). This indicates that the ability to become quiescent and to exit the quiescent state is not exclusive to the lab strain. Importantly, the ability to re-enter the cell cycle when the nutrient limitation was relieved was similar for single cells and cells with one or more buds (**Figure 4D**, **Figure 4E**, **Supplementary Figure 13A** and **Supplementary Figure 13B**). Thus, in *C. albicans,* quiescence is not unique to unbudded cells that are G1-arrested. We posit that the distinction between arrested budded and unbudded cells may be less critical for *C. albicans* survival of starvation than has been described for *S. cerevisiae*. Furthermore, the timing of reinitiation of division is dependent upon the nutrient composition of the media during the starvation period.

To address what contributes to the nutrient-dependent delay in the reinitiation of cell division following starvation, we tested the effect of glucose concentration, since it differed by 20-fold between the rich and minimal medium used. Specifically, we measured the dynamics of quiescence using SD-2%, which differs from minimal medium only in having 20-fold more glucose. Notably, cells starved in SD-2% glucose resumed cell division faster than cells starved in minimal medium, but slower than cells starved in rich medium (**Supplementary Figure 13C** and **Supplementary Figure 13D**). Thus, glucose concentration in the media used during starvation clearly affects the proportion of unbudded cells, and the rate at which they exit quiescence/resume division, and the other nutrients present in the rich medium likely contribute to the dynamics of quiescence as well.

### Quiescence reduces the cidality of antifungal drugs

Since quiescence in other organisms increases the ability of cells to tolerate a range of stresses (18,29), we compared the viability of proliferative and quiescent *C. albicans* cells following exposure to different fungicidal drugs. Cell viability was assessed using propidium iodide (PI) and SYTO9 staining quantified using flow cytometry (30,31) which we confirmed distinguishes live and dead proliferative and quiescent cells (**Supplementary Figure 14**). We first tested the viability of proliferative cells grown in rich media to increasing concentrations of micafungin, ranging from 0.001µg/mL to 10µg/mL using the lab strain (SC5314) and confirmed a dose dependent result (**Figure 5A**). Using the same drug concentrations we confirmed that quiescent cells also exhibited a dose-dependent increase in cell death. However, the loss in viability was dramatically higher in proliferative cells than in quiescent cells. For example, 0.1µg/mL of micafungin killed 90% of proliferative cells, whereas only 26% of quiescent cells were killed at this concentration. Moreover, at the maximum dose (10µg/mL) only 40% of quiescent cells were killed. We confirmed the differential effect of micafungin on proliferative and quiescent cells by plating cells following drug treatment and counting colony-forming units (CFUs) (**Figure 5A**).

**Figure 5.**
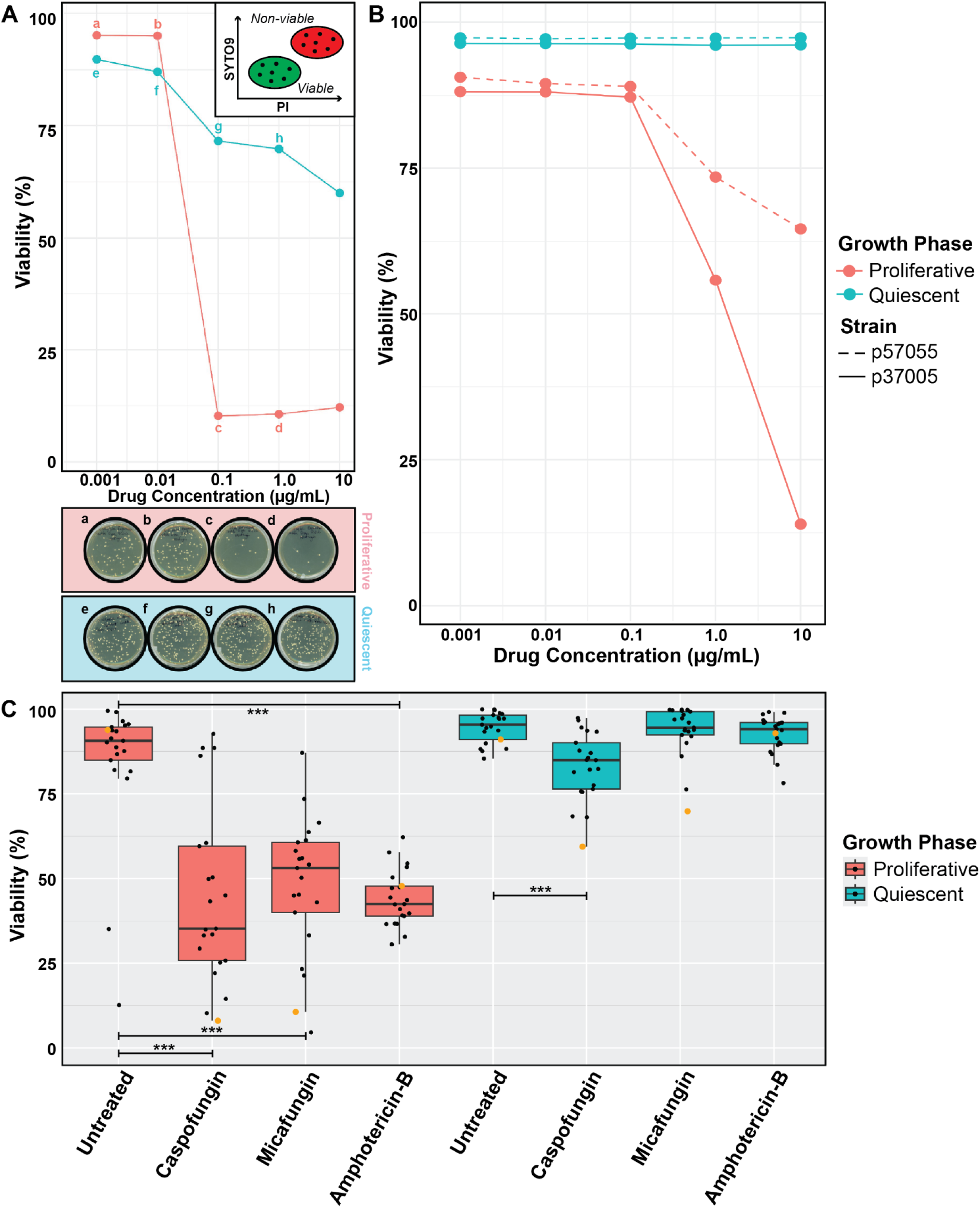
Quiescent cells exhibit increased survival when treated with antifungals. **A)** Viability of proliferative and quiescent cells using the lab strain SC5314 subjected to a range of doses from 0.001µg/mL to 10µg/mL of micafungin. Viability was determined using PI/SYTO9 staining and flow cytometry. Viability estimates of samples indicated with letters a-h were verified by plating ∼200 cells onto YPD plates and counting colony forming units (CFUs). **B)** Viability of proliferative and quiescent cells using the clinically isolated strains p57055 and p37005 at micafungin concentrations of 0.001 µg/mL to 10 µg/mL. **C)** Viability for proliferative and quiescent cells using 20 genetically diverse strains and the lab strain SC5314 at the LD50 drug concentration identified for proliferative SC5314 cells. Strains were treated with either 1µg/mL caspofungin, 1µg/mL micafungin, or 20µg/mL amphotericin B. Black line inside box shows median viability of 21 strains; box range indicates interquartile range (IQR); whiskers indicate 1.5 times the IQR from median. Viability of each strain is indicated as a black dot; SC5314 viability is indicated with a yellow dot; the p-value was determined by Student’s t-test, ∗p < 0.05, ∗∗p < 0.01, ∗∗∗p < 0.001.

To ask if this result is seen across different clinical isolates, we exposed two clinical strains, p57055 and p370005, to increasing amounts of micafungin. In proliferative cells, both strains exhibited increased drug resistance (**Figure 5B**) compared with the lab strain (**Figure 5A**). Remarkably, we observed no lethality (<1%) for both clinical strains when quiescent cells were treated with up to 10 µg/ml of micafungin (**Figure 5B**).

We then expanded the study to 20 genetically diverse strains and tested two additional antifungals: caspofungin and amphotericin B. Using the lab strain, we first confirmed increased cell death of proliferative cells with increasing drug concentrations of caspofungin (**Supplementary Figure 15**), cicafungin (**Supplementary Figure 16**), and amphotericin B (**Supplementary Figure 17**). By contrast, quiescent cells in the lab strain background displayed significantly higher viability when exposed to caspofungin (**Supplementary Figure 18),** micafungin (**Supplementary Figure 19**) and amphotericin B (**Supplementary Figure 20**). For each drug, we selected a concentration that resulted in killing at least 50% of proliferative cells in the lab strain. We assayed all 20 additional strains at this concentration for each of the three drugs in both proliferative and quiescent cells. Overall, clinical isolates had significantly decreased susceptibility relative to the lab strain for most of the drugs when treated as proliferative cells (**Figure 5C**). For all strains tested, quiescent cells had higher survival of drug treatment compared to proliferative cells (**Figure 5C**).

## Discussion

In this study we found that quiescence in *C. albicans* cells is characterized by physiological remodeling, increased cell density, activation of a unique gene expression program, and increased resilience to environmental and antifungal stresses. We found that the medium in which starvation and quiescence are established affects the degree of these physiological changes. These findings extend the understanding of fungal quiescence beyond model fungal systems, and reveal the potential clinical relevance of quiescence for pathogenic fungi.

Intracellular fluidity in quiescent cells differs between rich and minimal culture media. These two media also differ in pH, with minimal medium having pH 5.5 and rich medium having pH 6.5. In *S. cerevisiae,* lower pH *increases* the degree of cytoplasmic crowding and aggregate formation (27). However, in the *C. albicans* isolates studied here, *D*_eff_ was *higher* in the *lower-pH* minimal medium, relative to the *higher-pH* rich medium cultures. Precise measurements of intracellular pH will be needed to ask if cell medium pH causes the difference in cytoplasmic crowding observed.

The difference in initial glucose concentration between rich (2%) and minimal (0.1%) media is another factor that could contribute to the differences in quiescence and intracellular fluidity. *C. albicans* likely accumulates larger carbohydrate stores (i.e. trehalose and glycogen) in rich media, with additional trehalose uptake from the rich media (32). In *S. cerevisiae*, carbohydrate stores correlate with increased cellular density, enhance stress resistance during quiescence, and provide readily metabolizable energy upon quiescence exit (32).

Furthermore, the accumulation of trehalose and glycogen increases cellular viscosity and reduces diffusion in *S. cerevisiae* cells (33). We speculate that rich media promotes carbohydrate storage in quiescent *C. albicans* cells, whereas in minimal media less carbohydrate is stored due to less available glucose. Conversely, in minimal medium, the increased cytoplasmic fluidity in quiescent cells may result from decreased macromolecular content, particularly reduced ribosome and mRNA concentrations, due to reduced translational activity in quiescent cells growing in nutrient-poor conditions. This is consistent with the relative decrease in transcripts related to protein production in minimal media compared with rich media (**Figure 2B**). We posit that the nature of nutrient depletion shapes the physical properties of quiescence and, potentially, its protective capacity (34).

In *C. albicans,* quiescence confers increased survival to antifungal drug exposure. More quiescent cells than log-phase cells survived exposure to caspofungin, micafungin, and amphotericin B, which target active cellular processes including cell wall biosynthesis and membrane integrity (35). These results underscore the importance of considering the metabolic state of the cell when evaluating drug susceptibility, and point to quiescence as a likely contributor to treatment failure and recurrence in *Candida* infections.

Critically, *C. albicans* quiescent cells maintain the capacity to re-enter the cell cycle upon nutrient replenishment, fulfilling the definition of quiescence established for *S. cerevisiae* and *S. pombe* (36). This reversibility reinforces the idea that quiescence represents a bet-hedging strategy (28,37): an investment in long-term survival at the expense of short-term proliferation (38). This suggests that quiescent populations may act as reservoirs of infection (39), capable of withstanding immune or pharmacologic clearance and resuming growth when conditions permit. An open question is whether a subset of cells enter a quiescent state even when conditions are favorable, and thus contribute to phenotypic diversification of genetically homogeneous populations. Future studies should investigate the molecular regulators that govern quiescence entry and exit in *C. albicans*, particularly under *in vivo* conditions, and whether targeting these pathways can sensitize quiescent cells to antifungal agents. Recognizing and targeting quiescent fungal cells may be key to overcoming persistent infections and improving antifungal therapy outcomes.

## Materials and Methods

### Strains and media

The laboratory *C. albicans* strain SC5314 was used in all experiments unless otherwise indicated. Prior to all experiments, strains were streaked onto YPD plates supplemented with 2% glucose using sterile technique inside a laminar flow hood and parafilmed before being incubated for two nights in 30°C. A single *C. albicans* colony was used to inoculate a 5 mL culture in YPD and incubated overnight at 30°C with continuous rotation. 50 µL of the overnight culture was subsequently back-diluted into fresh 5 mL YPD and incubated for 6 hours prior to experimental setup. Cells are harvested by centrifugation at 4000 revolutions-per-minute (RPM) for 5 minutes to ensure removal of YPD medium, thereby preventing contamination of subsequent nutrient-limiting conditions. Cells are then resuspended in fresh YPD, 1X PBS or sterile HiPure water before getting counted with a hemocytometer. Cell number is adjusted per experimental conditions necessary.

All media were prepared in liquid form with pre-made solutions, sterilized by autoclaving, and supplemented with glucose post-cooling, depending on the media composition. Rich media (YPD) was formulated with 10 g Bacto-Yeast Extract [BD Diagnostic Systems, 212750] and 20 g Bacto-Peptone [ThermoFisher, 211677] and with MilliQ water being added to 1L. Minimal media was prepared by mixing 100 mL of 10X glucose salts with 896 mL MilliQ water, 1 mL 1000X concentrated metal solution, and 1 mL 1000X concentrated vitamin solution. The 10X glucose salts solution contained 1 g calcium chloride dihydrate, 1 g sodium chloride, 5 g magnesium sulfate heptahydrate, 10 g potassium phosphate monobasic, and 50 g ammonium sulfate, with MilliQ water added to 1 L. Minimal media was further supplemented with specific metal and vitamin mixtures prior to autoclaving. A 1000X concentrated metal stock solution contained 500 mg boric acid, 40 mg copper(II) sulfate pentahydrate, 100 mg potassium iodide, 200 mg iron(III) chloride hexahydrate, 400 mg manganese sulfate monohydrate, 200 mg sodium molybdate dihydrate, and 400 mg zinc sulfate heptahydrate. The 1000x concentrated vitamin stock solution included 2 mg biotin, 400 mg calcium pantothenate, 2 mg folic acid, 2 g inositol, 400 mg niacin, 200 mg p-aminobenzoic acid, 400 mg pyridoxine hydrochloride, 200 mg riboflavin, and 400 mg thiamine hydrochloride. The 10X glucose salts solution, the 1000X concentrated metal and vitamin solutions were pre-made to allow for the addition of 1 mL per liter of media during final preparation. Minimal media were tested using glucose concentrations of 5.6 mM, 2.8 mM, 1.4 mM, and 0.7 mM glucose concentrations were selected as the minimal media glucose concentration for all future experiments.

### Growth dynamics

Cells from an overnight pre-culture were counted and inoculated in a 96-well plate in 200µL for the duration indicated in each experimental condition. For the quantification of final yield at 72 hours, cells were incubated in the TECAN Plate Reader set at 30°C with no shaking. The starting OD was set to 0.1 for minimal conditions such that the final OD remained in the linear range (OD600 > 0.75) of plate reader. Absorbance was recorded from each well every 5 minutes, and the plate was incubated for 72 hours. To study the growth dynamics in rich media and to test if the final yield is dependent on glucose concentration, cells were incubated for 72 hours and diluted down to the linear range of the plate reader before the absorbance at 600nm was measured in triplicate for each condition. Growth curves were analyzed using the Growthcurver package in R (https://github.com/sprouffske/growthcurver). Outlier samples were removed before regression analysis and calculating R-square, significance and slope using the glm function in R.

### Bud index quantification

For the quantification of unbudded cells over time the following procedure was followed. Yeast cultures were incubated overnight in either minimal or rich media,the following day cells were counted with a hemocytometer and then diluted in the respective media to a final density of 1e5 cells/mL in 5mL glass tubes. The samples were incubated in a 30°C rotating incubator before a 50µL sample was collected at 2, 4, 6, 24, 48, and 72 hours post inoculation. The samples were diluted in sterile water in 1.5mL Eppendorf tubes at cell concentration in a microscope field-of-view had about 100 cells. Diluted cells were sonicated to ensure cells that have adhered to others were separated while mother cells that are actively budding remain connected to their daughter cells. The sonicated cells were transferred into a black 96-well plate with a flat glass bottom compatible with microscopy (Fisher Scientific, 165305) and imaged using a Nikon Ti2-E with the NIS Elements Acquisition Mode. During imaging, a number of images were acquired to make sure the total number of cells quantified exceeded 200 cells for each unique condition. The images acquired were exported as .TIFF files and analyzed using ImageJ. The ‘Cell Counter’ plugin was used to quantify the number of “Singlets” or unbudded cells, “Doublets” or budded cells, “Triplets” or a 3 cell cluster, and clusters with more than 3 cells as “Quadruplets”.

### Cell cycle analysis

A citrate buffer was prepared by mixing 14.71g of sodium citrate dihydrate and 1L of milliQ water (pH adjusted to 7.2 with citric acid monohydrate) and filter sterilized to prevent fermentation. Cells were assayed either after 6 hours post-inoculation for proliferative growth or 72 hours post-inoculation for quiescent growth profile. Each sample was individually collected, washed with 1X PBS, fixed with 80% ethanol, and stored at 4°C overnight.

The fixed samples were individually collected by centrifuging the cells at 6,000 x g for 1 minute at room temperature before being counted. Five million cells were used for staining per condition. The ethanol supernatant was removed and cells were resuspended in 1mL of citrate buffer. Samples were centrifuged, the supernatant was removed, and fresh 1mL of citrate buffer was added to perform a single wash. Subsequently, 200µL of a pre-made 250µg/mL RNase A solution (Fisher Scientific, 12091021) was added to each sample before incubation in a 50°C heat block for one hour. The incubated samples then received 20µL of 20mg/mL Proteinase K solution (Fisher Scientific, AM2548). The incubation durations for both RNase A and Proteinase K were optimized, as future staining optimization becomes highly challenging if the incubation time is reduced.

After incubation, samples were centrifuged and the pellet was resuspended in 1mL citrate buffer and washed twice. The washed samples were individually collected in 1mL citrate buffer and sonicated at the lowest power setting prior to staining and quantification. All samples were kept on ice following the sonication step until flow cytometry quantification.

Two hundred microliters of each sample solution was individually mixed with 200µL of 2µM SYTOX Green (Invitrogen, S7020). A Cytek Aurora Spectral Flow Cytometer was used to quantify the fluorescence spectrum of the forward and side scatter plots. Debris was excluded and cells were gated before the B2-A and forward scatter was visualized for each sample. The exported .fcs files were further quantified using CytoExploreR (https://dillonhammill.github.io/CytoExploreR/).

### Mitochondrial Imaging

For mitochondrial staining of proliferative cultures using Mitotracker Red FM (MT Red), 200 µL of mid-log culture from YPD and 1 mL of mid-log culture from SD 0.1% glucose were sampled and washed twice with PBS (without Ca²⁺ or Mg²⁺) at a 1:1 culture:PBS ratio. Cells were resuspended in a final volume of 100 µL in prewarmed (37°C) PBS. A prewarmed stock solution of 500 nM MT Red was added to the resuspended cultures to achieve a final concentration of 100 nM MT Red, and samples were incubated for 30 minutes at 37°C. For staining of quiescent cultures, 40 µL of 72-hour culture from YPD or SD 0.1% glucose were treated using the same protocol. Samples were immediately imaged at 100X magnification using 585 nm excitation and 630/75 nm emission (mCherry) to visualize mitochondria. Imaging of NAT-p.TDH3-mNeonGreen-SFC1 was performed using 470 nm excitation and 525/50 nm emission (GFP).

### Density fractionation

Density fractionation was performed using a discontinuous Percoll gradient. A Percoll working solution was prepared by diluting Percoll (Sigma-Aldrich) 9:1 (v/v) with 1.5 M NaCl to a final NaCl concentration of 167 mM. To generate the gradients, 10 mL of the Percoll solution was transferred into 15 mL centrifuge tubes and centrifuged at 13,800 RPM for 15 minutes at 20 °C using a fixed-angle rotor to establish a self-forming gradient.

Cell cultures were quantified by optical density measurements at 600 nm (OD₆₀₀), and 2 × 10⁹ cells (equivalent to 200 OD₆₀₀ units) were harvested by centrifugation. The cell pellet was resuspended in 1 mL of Tris buffer (15 mM Tris-HCl, pH 7.5) and carefully overlaid onto the preformed gradient. The gradients were centrifuged at 400 × g for 60 minutes at 20 °C in a tabletop centrifuge. Following centrifugation, visible fractions were imaged against a black background and image files are further quantified in ImageJ for the intensity changes of gray value across a clear area. An area is selected and the mean gray value is calculated across this area .

### RNAseq

Yeast cultures were grown in either rich or minimal media and sampled at multiple time points corresponding to 2, 5, 8, 11, 24, and 48 hours post-inoculation. Cell pellets were collected by centrifugation and immediately flash-frozen in liquid nitrogen for RNA extraction. Total RNA was extracted using a standard acid-phenol chloroform method, followed by DNase digestion to remove genomic DNA contamination. RNA integrity was assessed using TapeStation. Libraries were prepared from 1–2 ng of rRNA-depleted or poly(A)-enriched RNA using the Lexogen CORALL Total RNA-Seq Library Prep Kit (Cat. No. 095) according to the manufacturer’s protocol. First-strand cDNA was synthesized using random priming and strand displacement stop primers, followed by ligation of linker oligos containing unique molecular identifiers (UMIs) and partial Illumina adapters. Libraries were then purified using magnetic beads and reprecipitated to remove reaction components. Amplification and indexing were performed via PCR with Lexogen’s i7 single indices. Final libraries were purified and assessed for concentration and fragment size distribution using the Agilent Bioanalyzer and quantified by Qubit prior to sequencing with the NovaSeq 6000 Sequencing System.

The reads were basecalled using Picard IlluminaBasecallsToFastq version 2.23.8 with APPLY_EAMSS_FILTER set to false. Following basecalling, reads were demultiplexed using Pheniqs version 2.1.0. The entire process was executed using a custom nextflow pipeline, GENEFLOW (40,41). Raw FASTQ reads were subjected to quality control, including adapter trimming and filtering of low-quality reads using FastQC and Trimmomatic. Cleaned reads were aligned to the reference genome using the splice-aware aligner STAR, and read counts per gene were quantified using featureCounts to generate a raw count matrix.

Gene expression data were analyzed using the DESeq2 package in R. A generalized linear model was fitted using the formula ∼ Time + Media to account for both temporal and media-dependent effects on gene expression. Normalization and statistical testing were performed according to DESeq2 guidelines. Hierarchical clustering was used to group genes by temporal expression profiles. Genes with similar expression dynamics were subjected to Gene Ontology (GO) term enrichment analysis using OrgDB annotations sourced from UniProt and Candida Genome Database association files. Enrichment was evaluated for overrepresented biological processes using over-representation analysis. Over-represented GO term descriptions for each cluster were summarized using GPT-4o.

### Rheological probes

To image cytosolic and nuclear 40nm-GEMs in *C. albicans* cells, TIRF Nikon TI Eclipse microscope in highly inclined thin illumination mode (HILO) was used at 488 laser excitation with 100% power. Fluorescence was recorded with a scMOS camera (Zyla, Andor) with a 100x Phase, Nikon, oil NA = 1.4 objective lens (part number = MRD31901, pixel size: 0.093 μm). Cells were imaged at 100 Hz (10 ms per frame) for a total of 2 seconds.

The tracking of particles was performed with the Mosaic suite of FIJI, using the following parameters: radius = 3, cutoff = 0, Per/Abs = 0.05, a link range of 1, and a maximum displacement of 5 px, assuming Brownian dynamics.

All trajectories were then analyzed with the GEMspa (GEM single particle analysis) software package that we are developing in house (42): https://github.com/liamholtlab/GEMspa/releases/tag/v0.11-beta. Mean-square displacement (MSD) was calculated for every 2D trajectory, and trajectories continuously followed for more than 15 time points were used to fit with linear time dependence based on the first 10 time intervals to quantify time-averaged MSD: MSD(*T*) = 4*D*_eff_*T*, where *T* is the imaging time interval and *D*_eff_ is the effective diffusivity with the unit of μm^2^/s. To determine the ensemble-time-averaged mean-square displacement (MSD), all trajectories were fitted with MSD(*τ*)_T−ens_=4*Dτ^α^* where *α* is the anomalous exponent, with *α* = 1 being Brownian motion, *α* < 1 suggests sub-diffusive motion and *α* > 1 as super-diffusive motion.

To generate region of interest (ROI) for marking individual yeast cell, cellpose python package (https://github.com/mouseland/cellpose) was used to segment based on the bright field image.These ROI were then input into GEMspa to quantify single cell effective diffusion, with at least three trajectories averaged for individual yeast cells. We used the median value of *D*_eff_ for single cell data to represent each condition. We measured the ROI area to deduce the cell volume based on 2D bright field images, assuming individual yeast cell are ellipsoidal, the Vcell = 4/3*π*r^3^.

### Microcolony assay

A 96-well glass-bottom plate was pre-treated for 30 minutes with 45 µL of 1 mg/mL Concanavalin A (ConA) to affix cells in place. The ConA was removed and wells were washed twice with sterile double-distilled water, then dried in a laminar hood during cell preparation. Cell cultures were grown in 5 mL glass tubes with continuous rotation in either YPD or SD 0.1% glucose media. Cells were sampled at mid-log phase (proliferative) and at 72 hours (quiescent) and gently washed twice with PBS without Ca²⁺ or Mg²⁺. Optical density was measured at 600 nm and cultures were diluted in double-distilled water to an OD₆₀₀ of 0.05. Five microliters of the diluted cell culture were added to 50 µL fresh YPD with thorough pipetting.The 96-well glass-bottom plate containing cell culture was centrifuged at 200 × g for 5 minutes on top of a clean KimWipe to prevent dust and scratches on the glass bottom. Using an inverted Nikon Eclipse Ti microscope with a 40× objective and NIS Elements Software, plate coordinates were selected and imaged every 15 minutes for 9 hours at 30°C. Cell morphology and time of bud emergence were manually analyzed using NIS Elements Software.

### Dose response assay

Twenty distinct *Candida albicans* strains (43) were inoculated into 5 mL of rich media in sterile 15 mL tubes and incubated overnight at 30°C with rotation. Each strain was cultured from a single colony under sterile conditions in a biosafety cabinet pre-cleaned with 80% ethanol and 10% bleach. Stock solutions of 5 mg/mL caspofungin, micafungin, and amphotericin B were prepared according to manufacturer specifications. A five-step serial dilution was performed using DMSO in a fume hood to reach final drug concentrations of untreated, 0.001 μg/mL, 0.01 μg/mL, 0.1 μg/mL, 1 μg/mL, and 10 μg/mL in the appropriate wells. Each intermediate dilution was generated by adding 100 μL of the higher concentration to 900 μL of DMSO, with thorough vortexing between steps. Aliquots were stored at −20°C and used within one month to minimize degradation.

For exponential phase assays, ^6^ cells/mL were counted and aliquoted within 1mL of rich media (YPD) into 1.5mL Eppendorf tubes and incubated at 30°C with shaking for 6 hours before getting treated. For quiescent phase assays, each strain’s culture was directly added into wells after 72 hours of incubation at 30°C with shaking. A single 2mm glass bead was placed in each tube to facilitate agitation of the culture walls during incubation avoiding cells sticking onto surfaces. After 24 hours, 100 μL aliquots from all samples were transferred into a 5mL round bottom polystyrene tube containing 400 μL of UltraPure water for downstream viability assessment. Samples were gently mixed by vortexing and subjected to flow cytometric analysis using a Cytek instrument to determine drug response across the strain and dosage series.

## Data Availability

Sequencing data is available at SRA PRJNA1271226. Scripts for all other analysis and figures, as well as all data, are also available on Github: https://github.com/GreshamLab/Quiescence_Candida_albicans.

## Acknowledgements

We are grateful for funding from NIGMS (R35GM153419), NSF (1818234), and BSF (2021276) (DG). We acknowledge the Zegar Family Foundation for their generous support. This project also received funding from the European Research Council (ERC) under the European Union’s Horizon 2020 research and innovation programme (grant agreement No 951475) (JB).

## Supplementary tables

**Table S1.** *Candida albicans* strains used in this study. The provenance of each strain is indicated.

| Strain Name | Origin | Geographical Location |
| --- | --- | --- |
| SC5314 | ATCC | Unknown |
| SC5314, pRPS10-40nm-GEMs (CytoGEMs) | Holt lab (LH4903) | Unknown |
| SC5314, pERG13-NLS-40nm-GEMs (NucGEMs), Nup49-mScarlet | Holt lab (LH4905) | Unknown |
| Sc5314, NAT-pTDH3-mNeonGreen-SFC1 | Berman Lab (CB1759) | Unknown |
| GC75 | Healthy individual;<br>oral sample | South Africa |
| p75010 | Bloodstream isolate | Belgium |
| p87 | HIV+ patient;<br>oral sample | South Africa |
| p37039 | Bloodstream isolate | New Jersey, USA |
| p75016 | Bloodstream isolate | Israel |
| p60002 | Bloodstream isolate | Arizona, USA |
| p78042 | Bloodstream isolate | Indianapolis, USA |
| p78048 | Bloodstream isolate | Manitoba, Canada |
| p75063 | Bloodstream isolate | France |
| p94015 | Bloodstream isolate | Utah, USA |
| p34048 | Bloodstream isolate | Turkey |
| p76055 | Bloodstream isolate | Iowa, USA |
| 12C | Vaginitis patient;<br>oral sample | Michigan, USA |
| L26 | Vaginitis patient;<br>vaginal sample | Iowa, USA |
| p57072 | Bloodstream isolate | Iowa, USA |
| p57055 | Bloodstream isolate | Nebraska, USA |
| p37037 | Healthy individual;<br>oral sample | Wisconsin, USA |
| p76067 | Bloodstream isolate | Ontario, Canada |
| 19F | Vaginitis patient;<br>vaginal sample | Michigan, Canada |

## Supplementary Figures

**Supplementary Figure 1.**
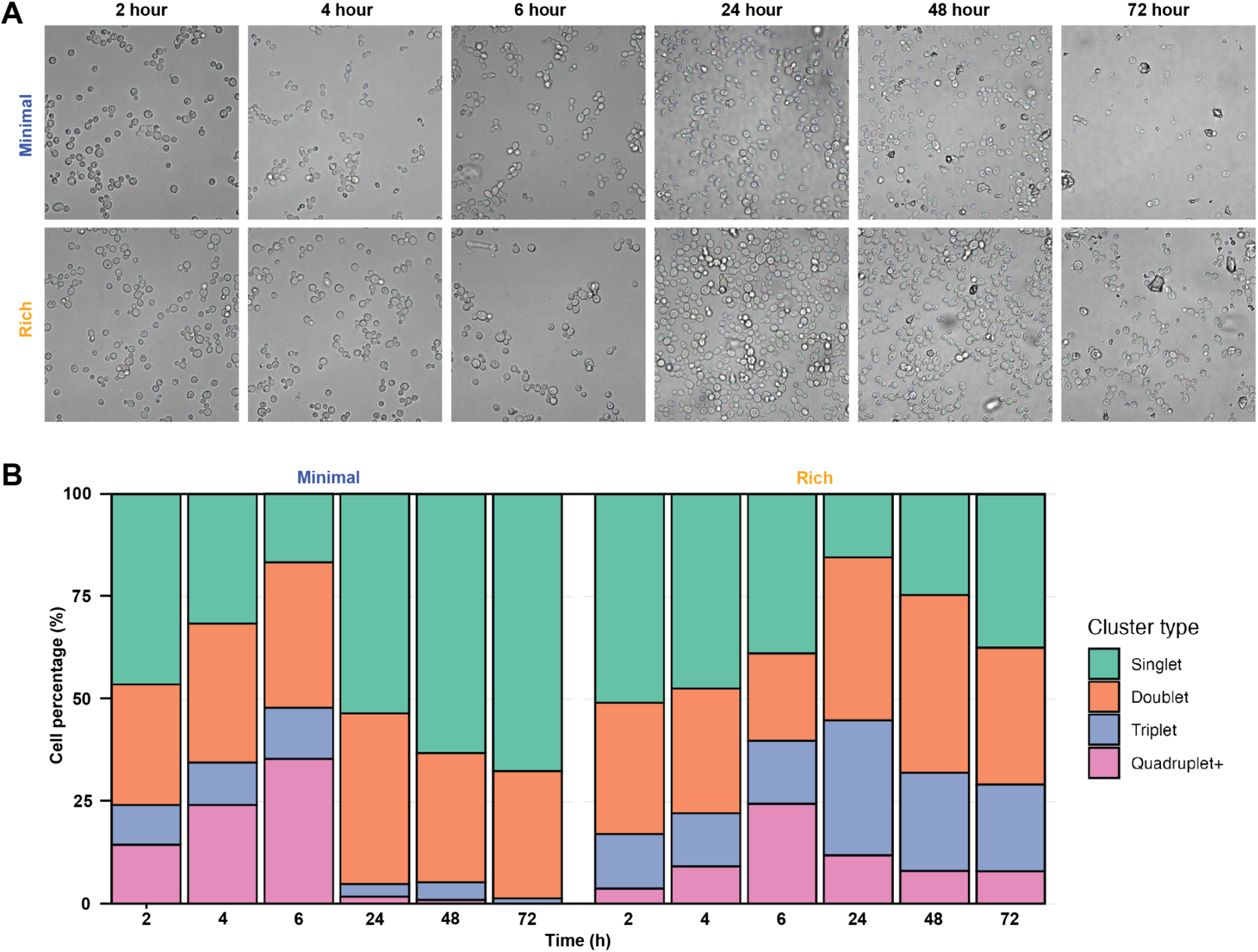
Phase contrast microscopy images of rich and minimal media-grown proliferative and quiescent cells used for bud index quantification. **A)** Representative phase contrast microscopy images showing cells at different growth phases grown in rich and minimal media. Cells are counted and marked as 1 (singlet), 2 (doublet), 3 (triplet), 4 (quadruplet+) or 5 (ambiguous) and counted using Cell Counter plugin for ImageJ. **B)** Cell cluster type percentages change with time in cells grown in rich and minimal media. Counts quantified from Cell Counter outputs; minimum of 200 cells were counted per condition.

**Supplementary Figure 2.**
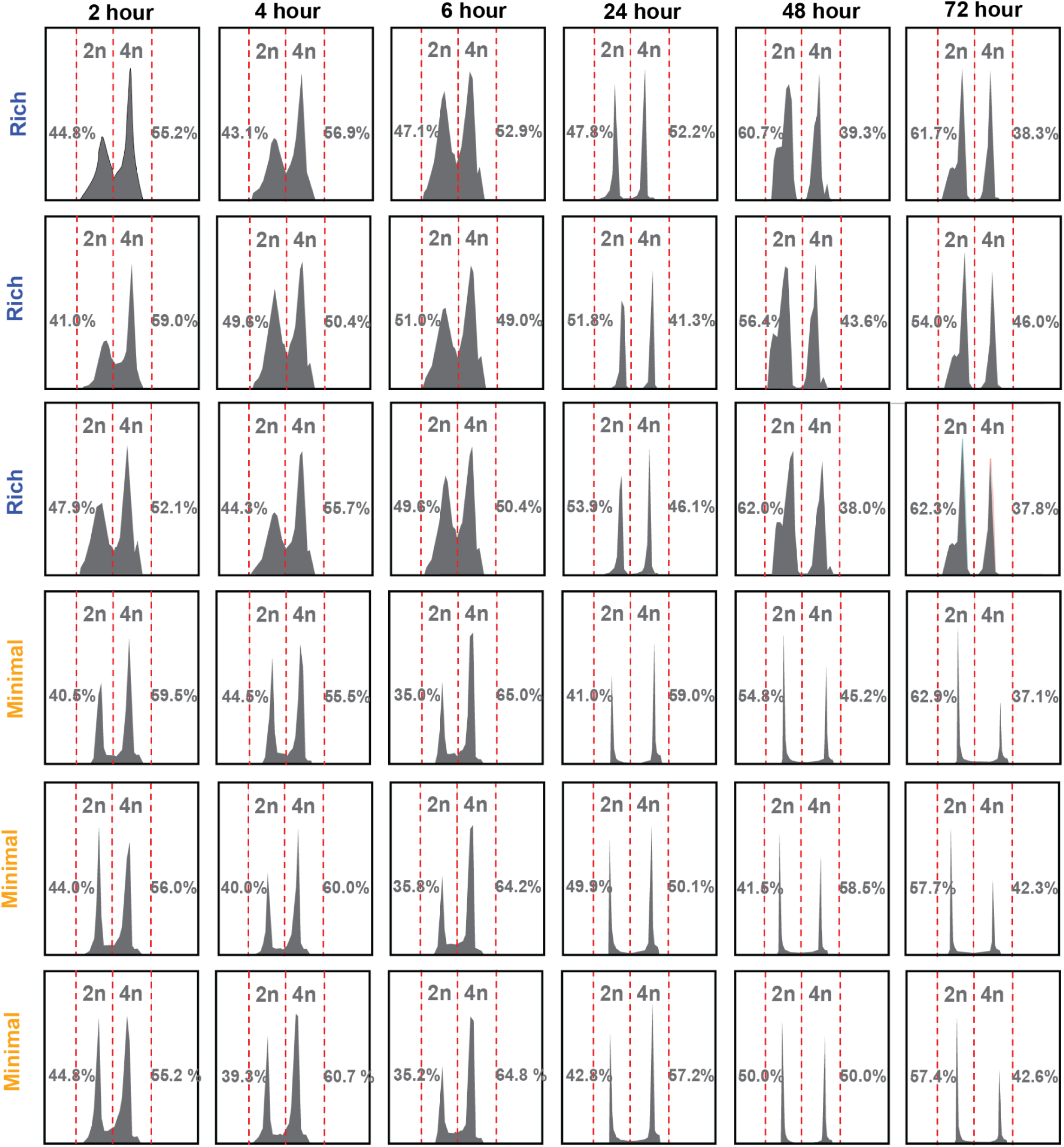
DNA content analysis of proliferative and quiescent cultures in rich and minimal media. Distribution of 2n and 4n DNA peaks. N = 50,000 events for each panel.

**Supplementary Figure 3.**
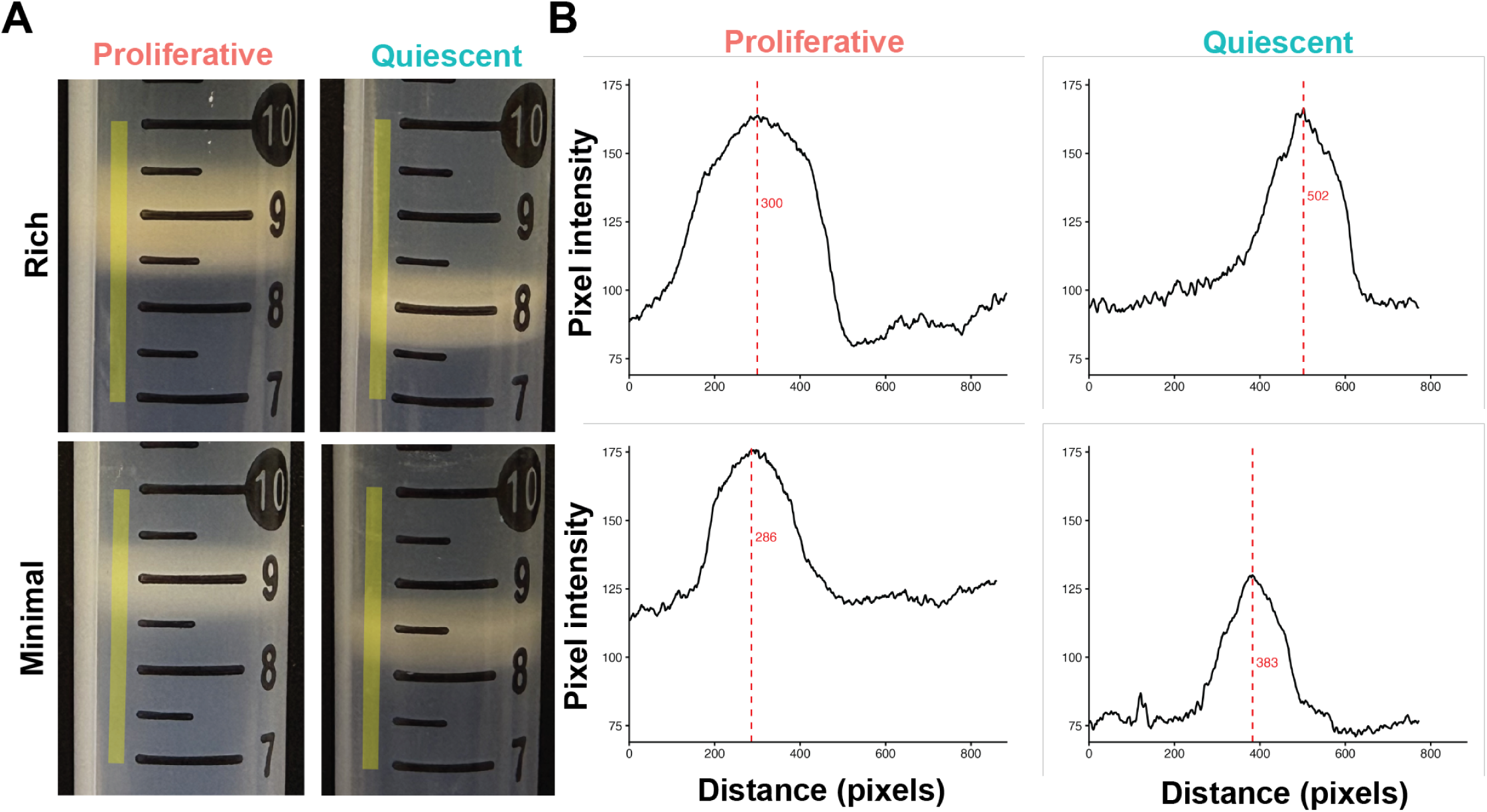
Density fractionation of proliferative and quiescent cultures in rich and minimal media. **A)** Images of density fractionation indicating the region assessed for pixel quantification(yellow vertical line). **B)** Pixel intensity distribution across the analyzed region with maximum distance at which peak pixel intensity was achieved is indicated (red line with red text).

**Supplementary Figure 4.**
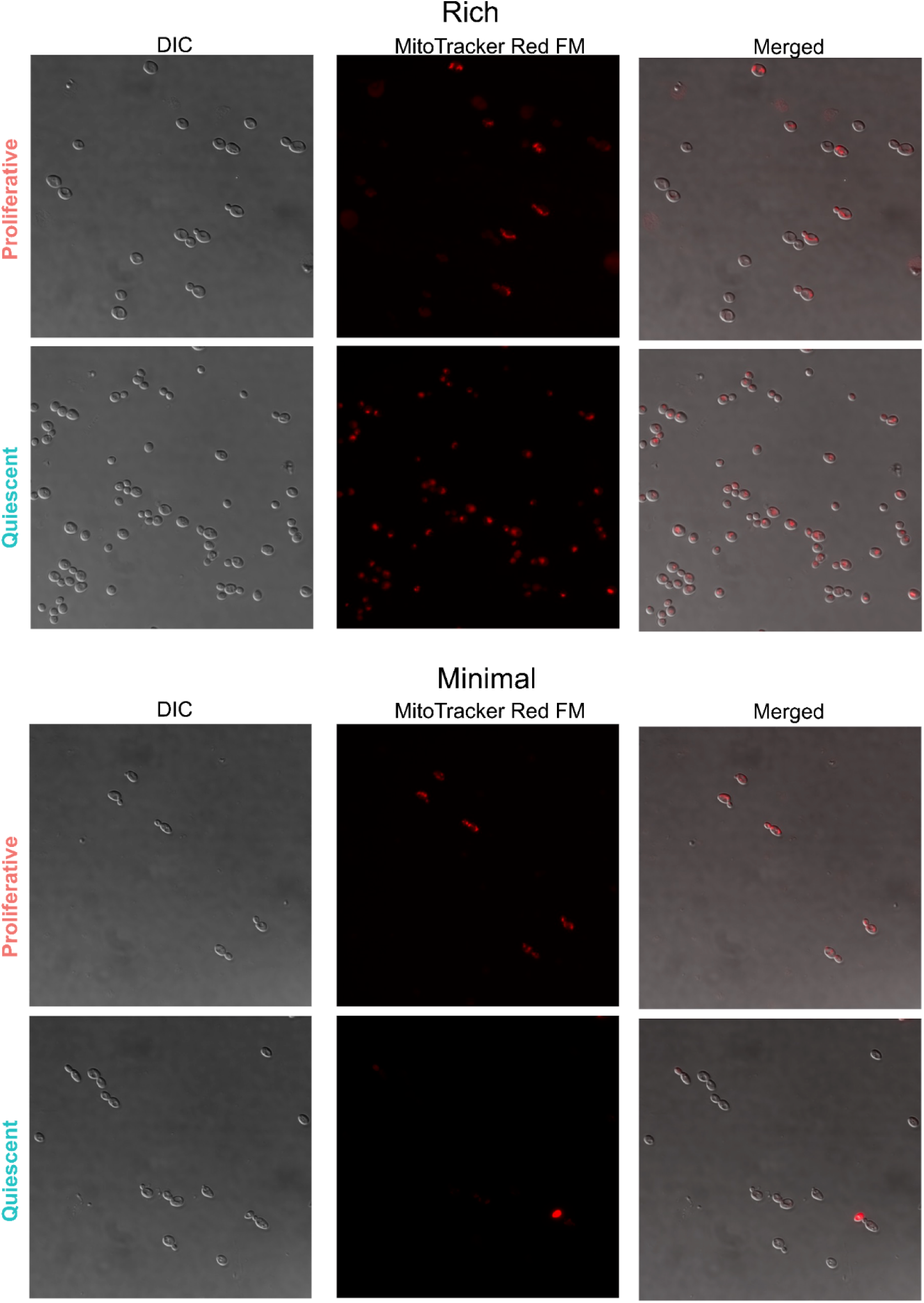
Mitotracker Red FM staining of proliferative cultures in rich media. SC5314 was stained with 100 nM MT Red for 30 minutes at 37°C. MT Red fails to selectively stain mitochondria of cells from rich and minimal quiescent cultures.

**Supplementary Figure 5.**
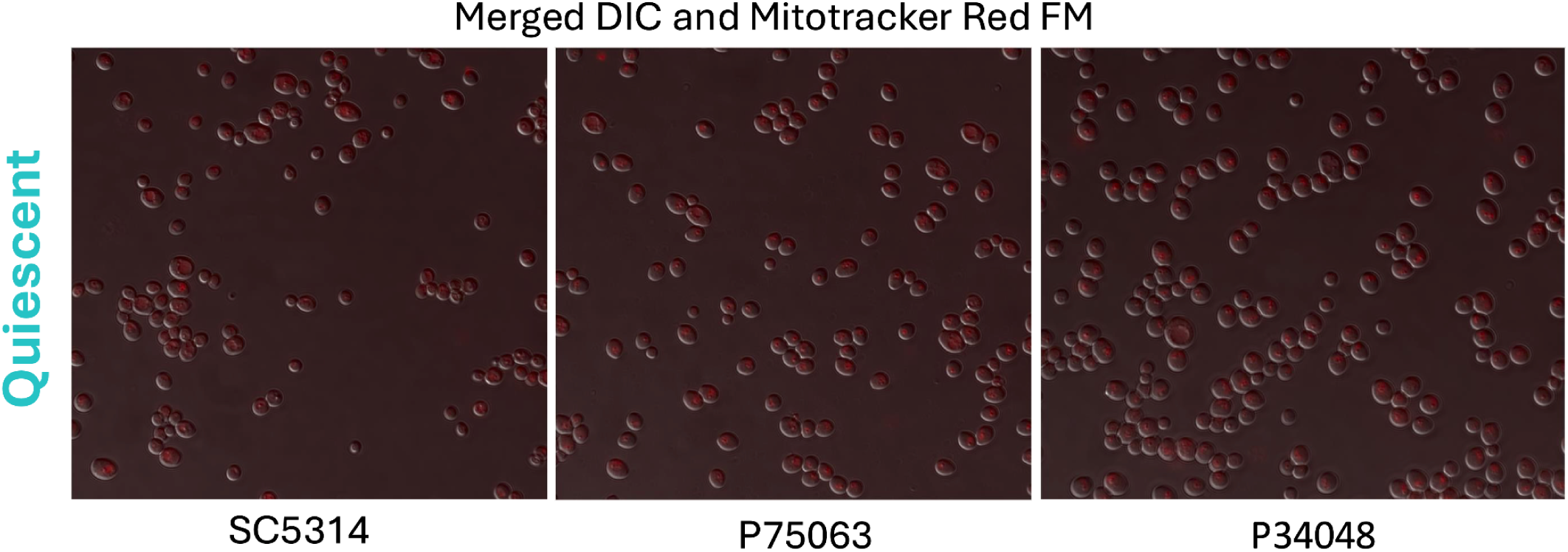
Mitotracker Red FM fails to selectively stain mitochondria in quiescent cultures of the lab strain SC5314 and two clinical isolates. Quiescent cultures of SC5314, P75063, and P34048 grown in rich media, incubated with Mitotracker Red FM, and imaged.

**Supplementary Figure 6.**
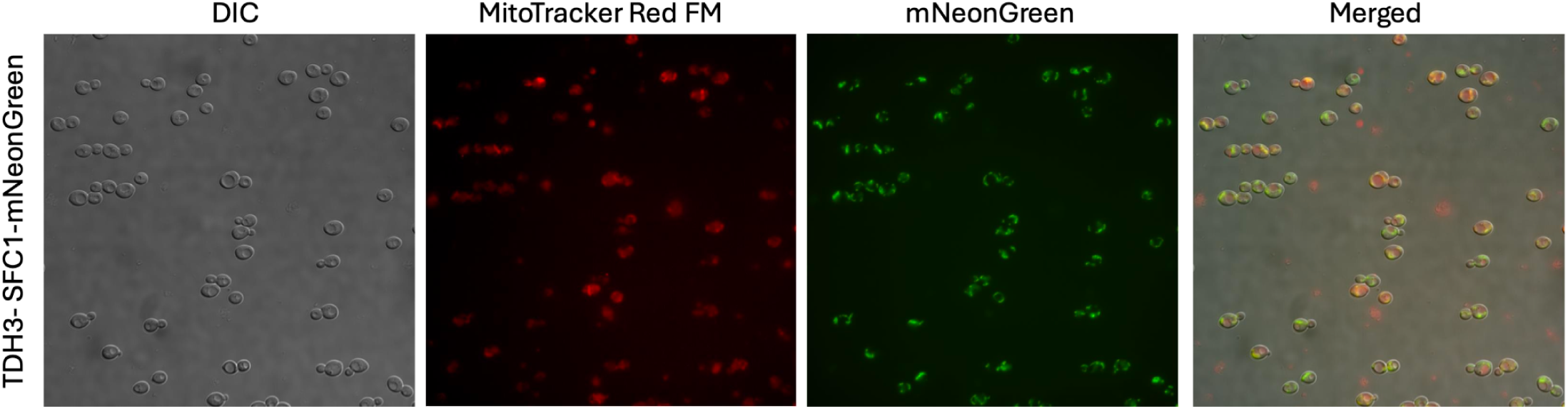
Colocalization of mNeonGreen-SFC1 and Mitotracker Red FM. Proliferative cells containing the mNeonGreen-SFC1 protein fusion were grown in rich media and stained with Mitotracker Red FM.

**Supplementary Figure 7.**
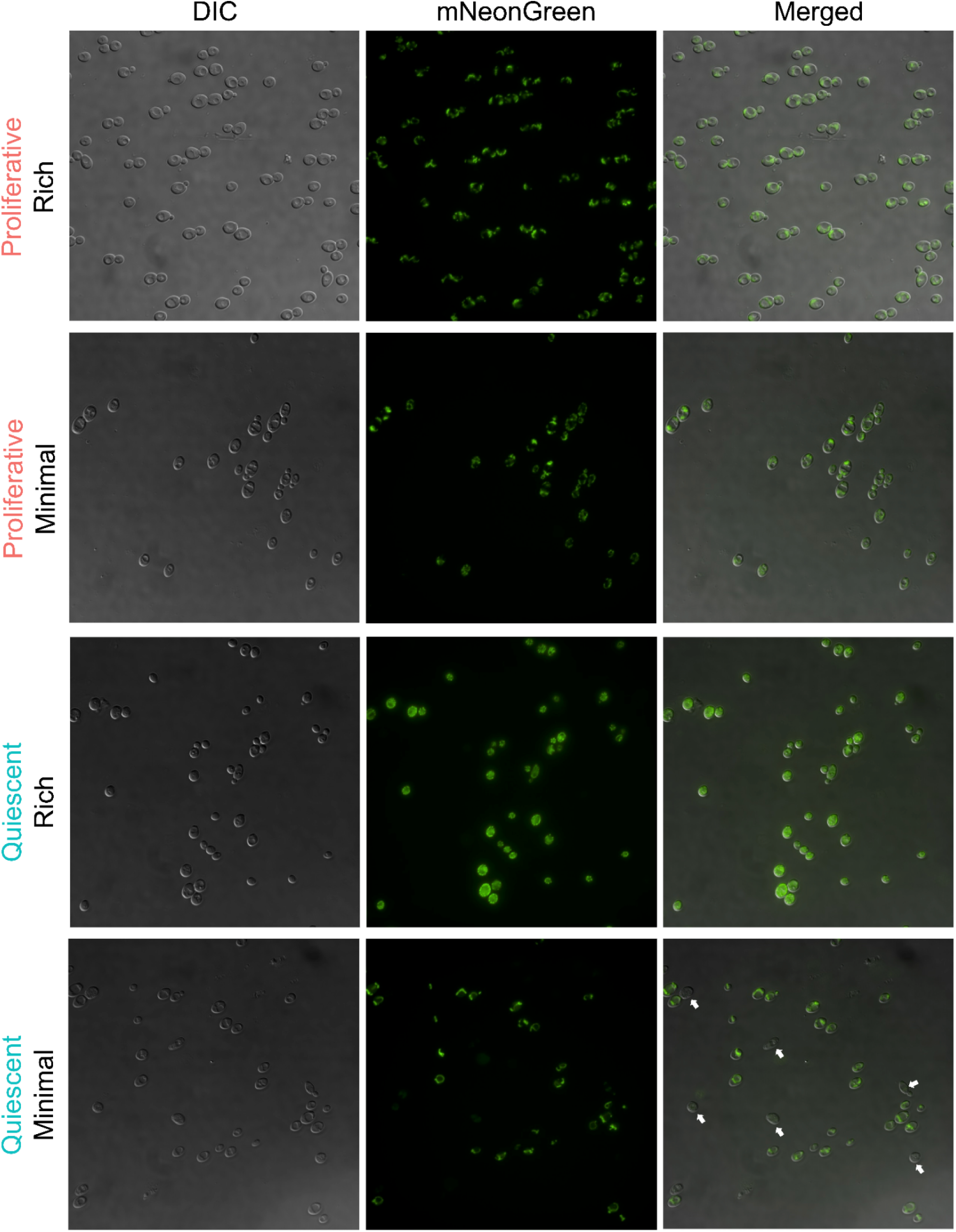
Imaging of mitochondria using mNeonGreen-SFC1 in quiescent and proliferative cells grown in rich and minimal media. White arrows indicate cells that lack any mNeonGreen-SFC1 signal.

**Supplementary Figure 8.**
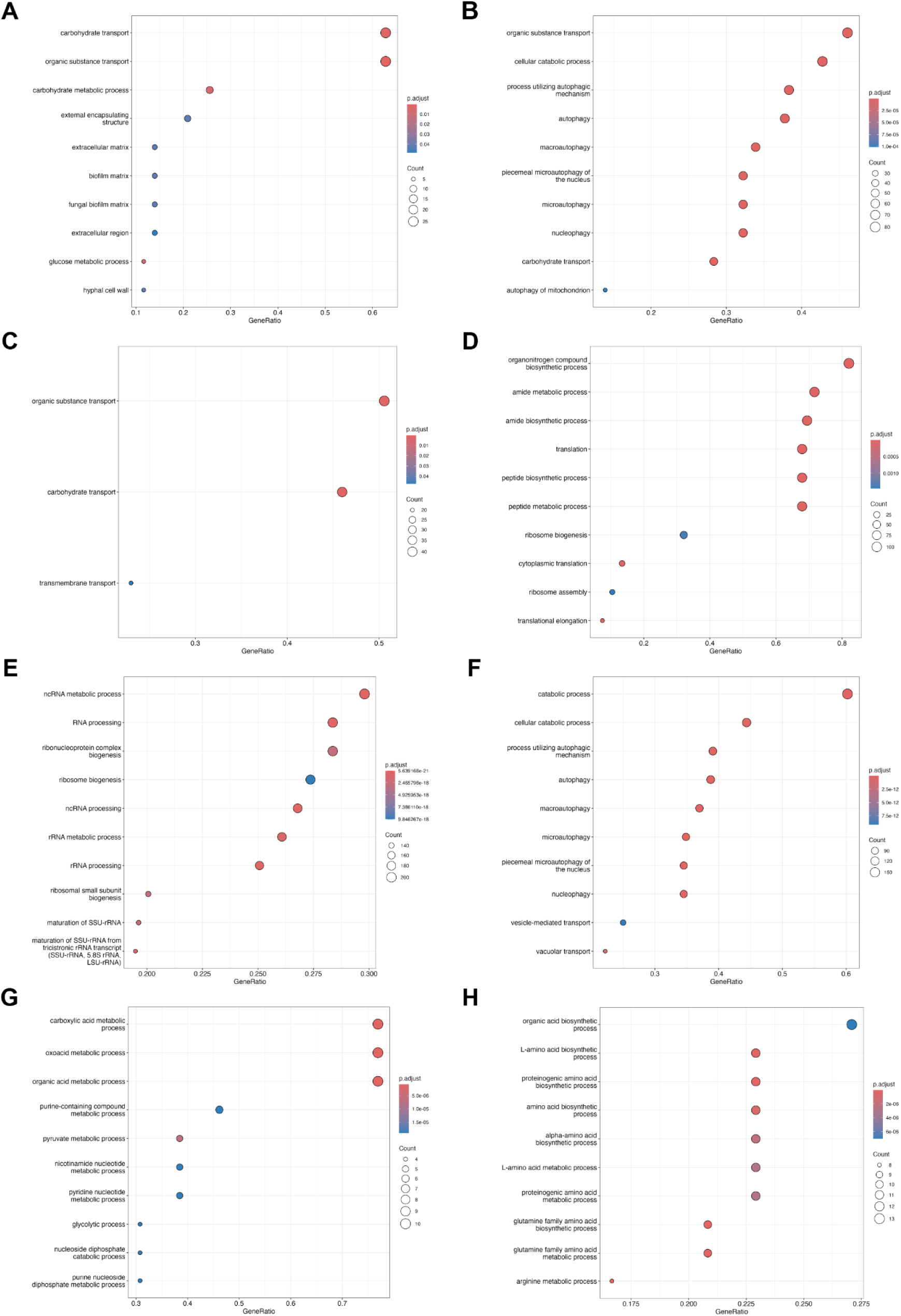
Overrepresentation analysis of differentially expressed gene clusters defined in Figure 2. Dotplot showing over–represented GO term descriptions for genes in **A)** cluster 1, **B)** cluster 2, **C)** cluster 3, **D)** cluster 4, **E)** cluster 5, **F)** cluster 6, **G)** cluster 7, and **H)** cluster 8.

**Supplementary Figure 9.**
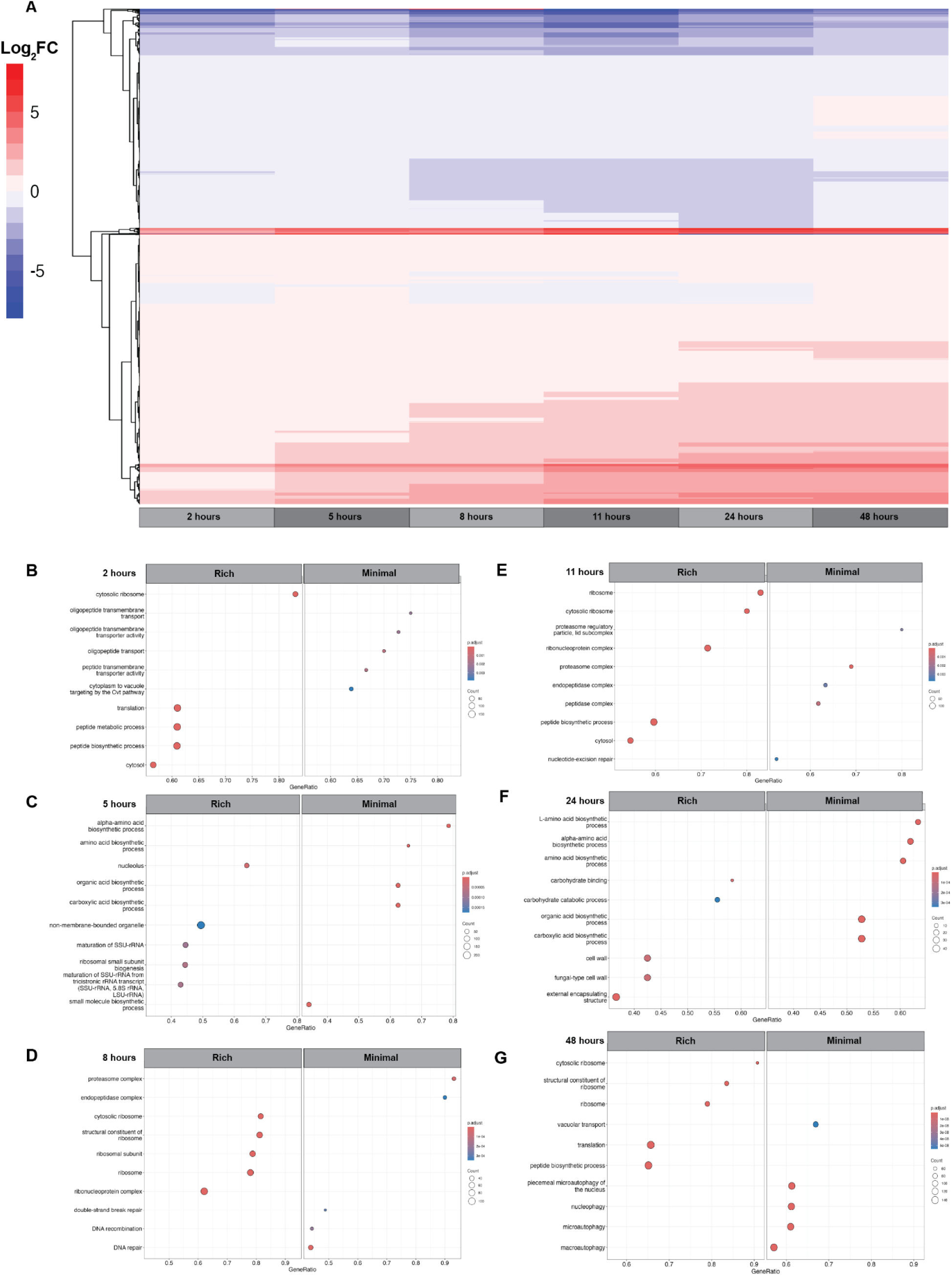
Differential gene expression analysis of media conditions. **A)** Heatmap of the relative log_2_-transformed fold changes of gene expression in rich media compared to minimal media at each time point. The data were clustered using hierarchical clustering. Gene Set Enrichment Analysis (GSEA) was used to find enriched GO terms at **B)** 2 hours post-inoculation, **C)** 5 hours post-inoculation, **D)** 8 hours post-inoculation, **E)** 11 hours post-inoculation, **F)** 24 hours post-inoculation, and **G)** 48 hours post-inoculation. Only significant GO terms (adjusted p-value < 0.05) are shown.

**Supplementary Figure 10.**
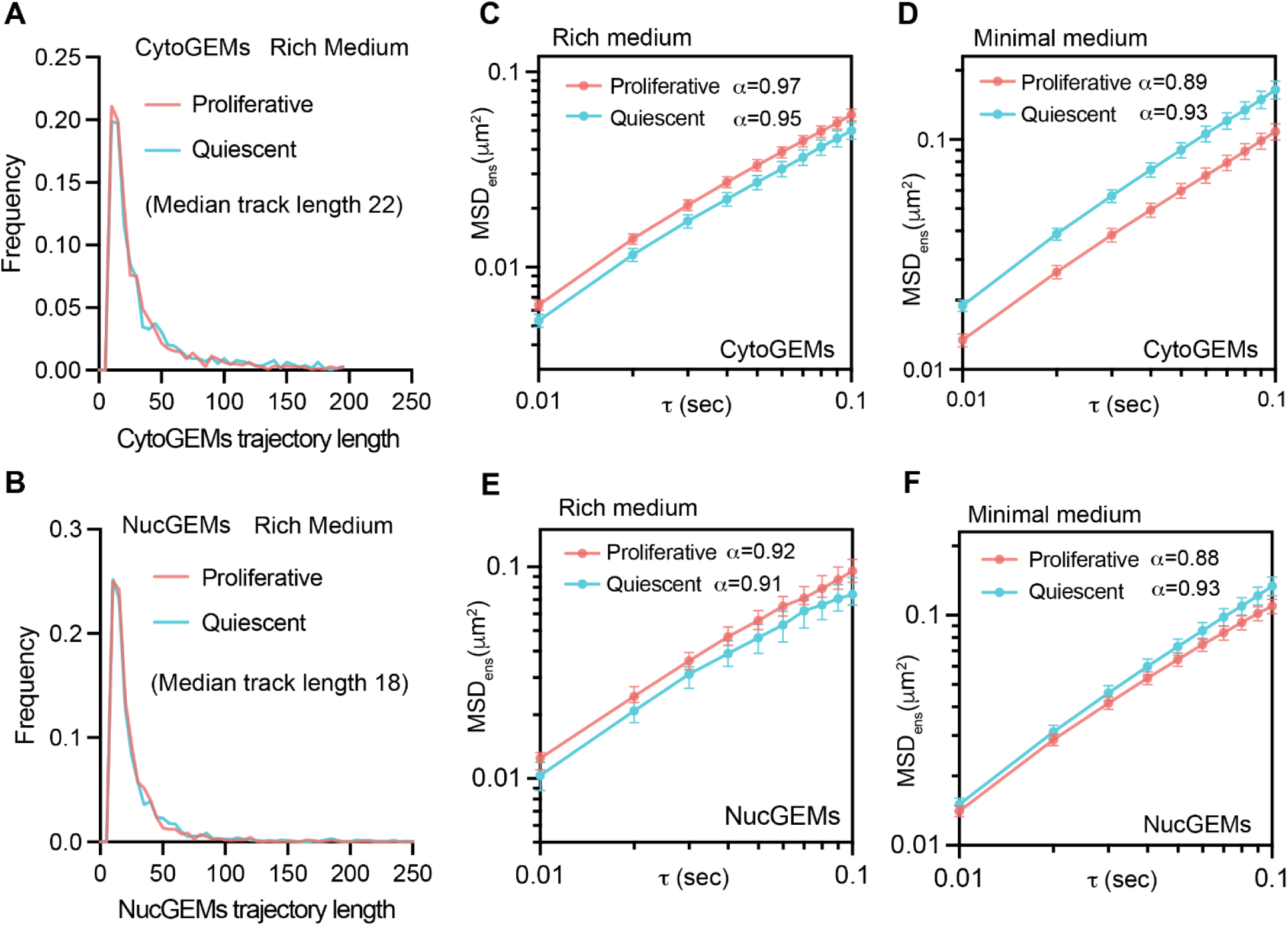
Molecular Crowding Profiles of SC5314 Nucleus and Cytoplasm in Rich and Minimal Media. **A)** The frequency distribution of cytoplasmic 40nm-GEMs trajectory length in rich media. Proliferative: n = 1434 trajectories; quiescent: n = 1280 trajectories. **B)** The frequency distribution of nuclear 40nm-GEMs trajectory length in rich media. Proliferative: n = 1570 trajectories; quiescent: n = 1567 trajectories. **C)** Ensemble-averaged mean-squared displacement (MSD) versus time delay (τ), log_10_ scale, for cytoplasmic GEMs in cells growing in rich medium. Proliferative: n = 111 cells; quiescent: n = 80 cells. **D)** MSD vs. τ plot for cytoplasmic GEMs in cells growing in minimal medium. Proliferative: 114 cells; quiescent: 58 cells. **E)** MSD vs. τ for nuclear GEMs in cells growing in rich medium. Proliferative: n = 139 cells; quiescent: n = 150 cells. **F)** MSD vs. τ for cytoplasmic GEMs in cells growing in rich medium. Proliferative: n = 140 cells; quiescent: n = 136 cells. For the MSD quantifications, a linear model was fit to determine the anomalous exponent α values for 40nm-GEMs in *C. albicans* cells. Median ± 95% confidence interval.

**Supplementary Figure 11.**
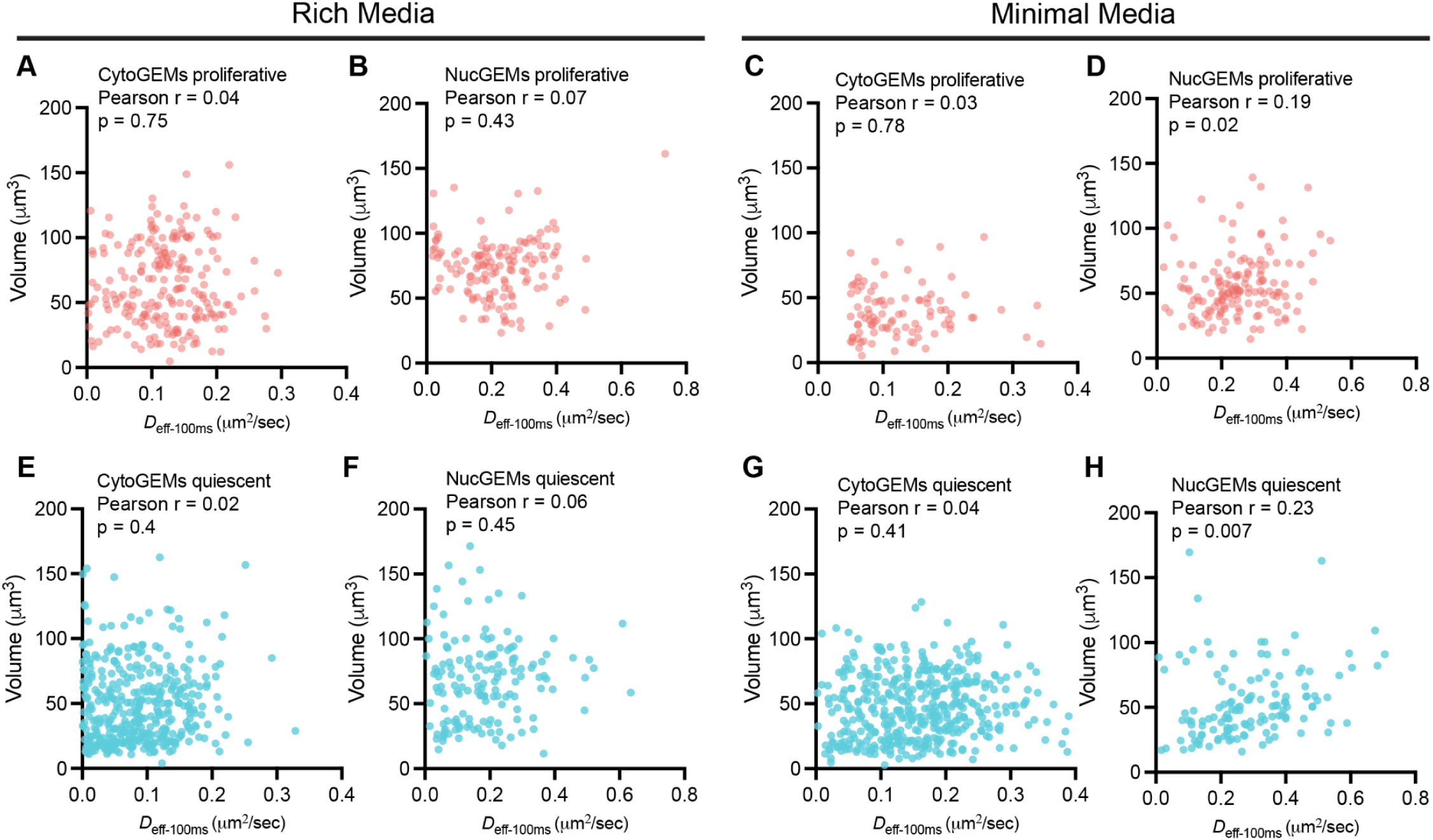
Relationship between cell volume and 40nm-GEMs effective diffusion coefficients. **A)** Cell volume to 40nm-GEM diffusion coefficient scatterplot for GEMs located in the cytoplasm of proliferative cells grown in rich media. **B)** Cell volume to 40nm-GEM diffusion coefficient scatterplot for GEMs located in the nucleus of proliferative cells grown in rich media. **C)** Cell volume to 40nm-GEM diffusion coefficient scatterplot for GEMs located in the cytoplasm of proliferative cells grown in minimal media. **D)** Cell volume to 40nm-GEM diffusion coefficient scatterplot for GEMs located in the cytoplasm of proliferative cells grown in minimal media. **E)** Cell volume to 40nm-GEM diffusion coefficient scatterplot for GEMs located in the cytoplasm of quiescent cells grown in rich media. **F)** Cell volume to 40nm-GEM diffusion coefficient scatterplot for GEMs located in the nucleus of quiescent cells grown in rich media. **G)** Cell volume to 40nm-GEM diffusion coefficient scatterplot for GEMs located in the cytoplasm of quiescent cells grown in minimal media. **H)** Cell volume to 40nm-GEM diffusion coefficient scatterplot for GEMs located in the nucleus of quiescent cells grown in minimal media. Pearson correlation r is calculated between diffusion coefficient (x-axis) and cell volume (y-axis) (Each dot represents one cell, p value was calculated by two tailed t test).

**Supplementary Figure 12.**
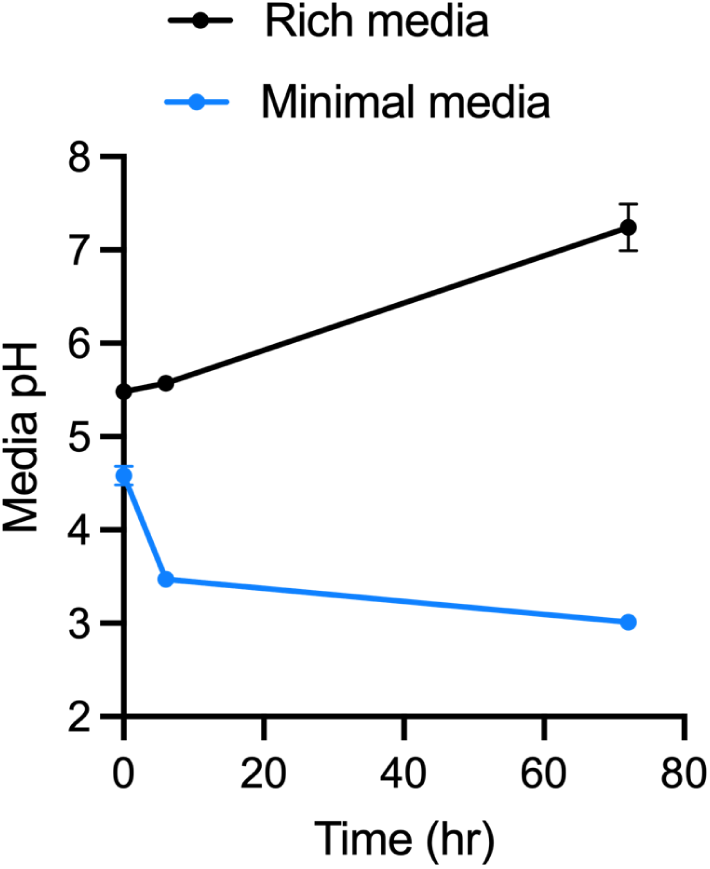
Culture media pH was measured at the indicated time points after C. albicans growth. Three culture tubes were used in two types of media (mean ± standard deviation)

**Supplementary Figure 13.**
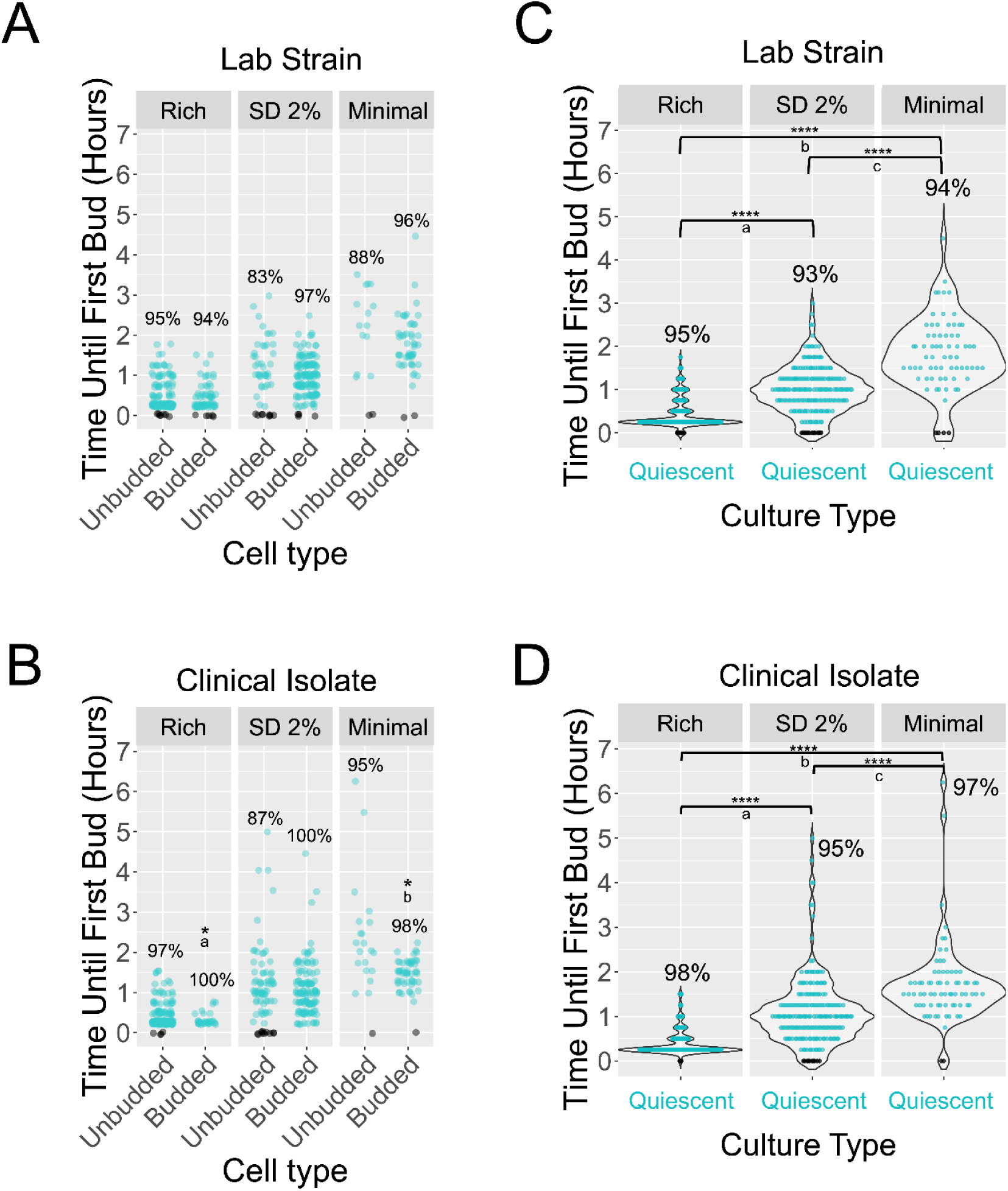
Time to bud emergence and budding status of the lab strain and a clinical isolate. **A)** Comparison of unbudded versus budded time until first bud emergence in quiescent cultures of the lab strain SC5314 in rich, SD 2% glucose, and minimal medium. The percentage of cells that divided within the 9-hour timeframe is noted above each bud category. **B)** Comparison of unbudded versus budded time until first bud emergence in quiescent cultures of the clinical isolate P34048 in rich, SD 2% glucose, and minimal medium. The percentage of cells that divided within the 9-hour timeframe is noted above each bud category. p-values for: a) 0.0437; b) 0.0105. **C)** Time until first bud emergence from quiescent cultures of the lab strain SC5314 grown in three types of media (rich and minimal media show the same data from Figure 4B). p-values for a) 2.40E-29; b) 6.51E-22; c) 2.08E-11 **D)** Time until first bud emergence from quiescent cultures of the clinical isolate P34048 grown in three types of media (rich and minimal media show the same data from Figure 4B). p-values for a) 4.08E-28; b) 6.80E-20; c) 1.22E-06. SC5314 rich quiescent, n = 243; SC5314 SD 2% quiescent, n = 199; SC5314 minimal quiescent, n = 70; P34048 rich quiescent, n = 208; P34048 SD 2% quiescent, n = 205; P34048 minimal quiescent, n = 74. p-values for significant differences between conditions or budding status are denoted with asterisks. The p-value was determined using Student’s t-test, ∗p < 0.05, ∗∗p < 0.01, ∗∗∗p < 0.001, ∗∗∗∗p < 0.0001).

**Supplementary Figure 14.**
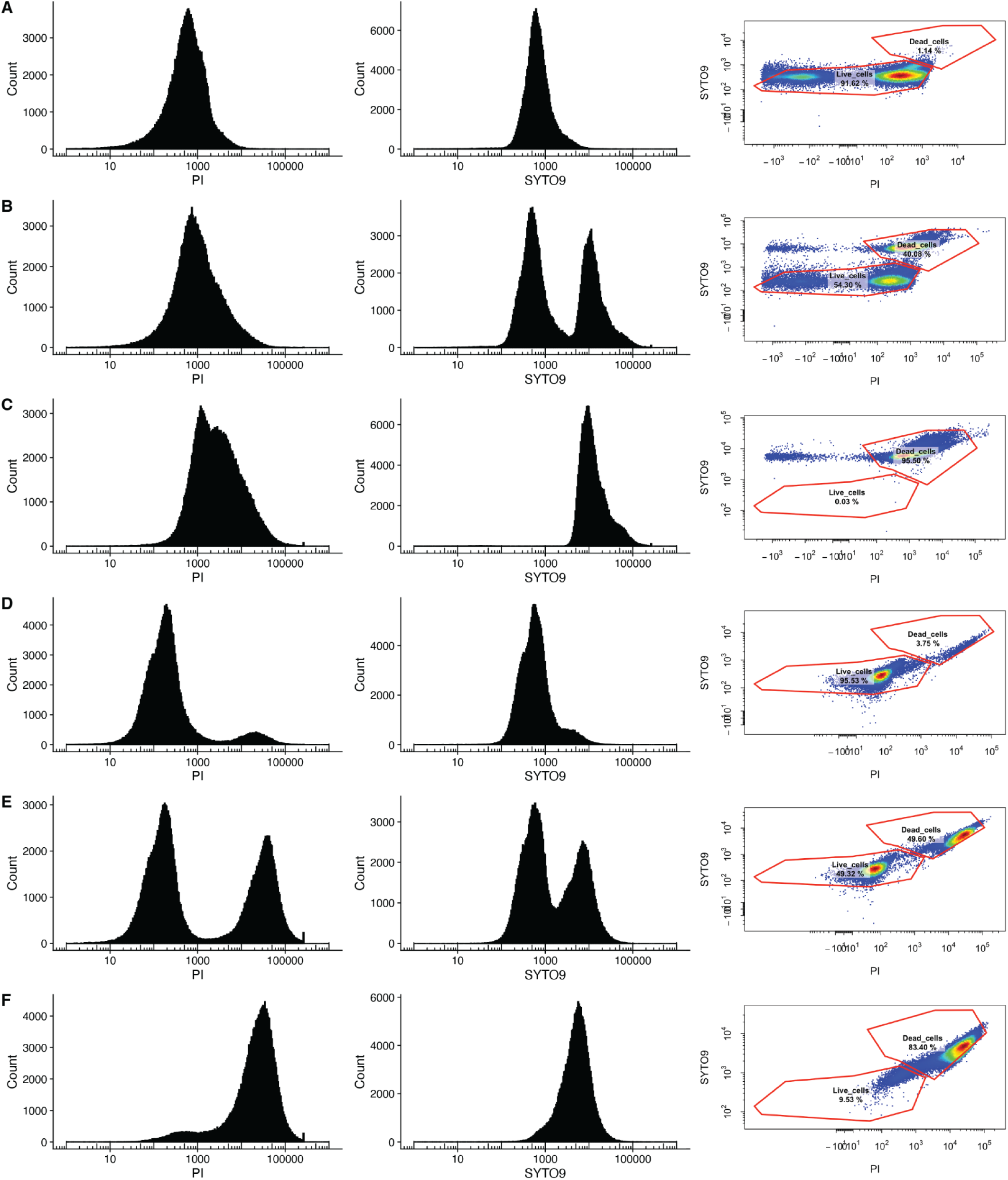
Validation of PI/SYTO9 staining and flow cytometry to measure viability of proliferative and quiescent cells. In each panel the distribution of PI (left panel), SYTO9 (middle panel) and the scatter plot of PI versus SYTO9 used to define flow cytometry gates is displayed for **A)** untreated proliferative cells grown (100% viability expected), **B)** a 50:50 mixture of untreated cells and cells exposed to 68℃ for 1 hour (50% viability expected) **C)** cells exposed to 68℃ for 1 hour (100% inviability expected) **D)** untreated quiescent cells (100% viability expected), **E)** a 50:50 mixture of untreated quiescent cells and quiescent cells exposed to 68℃ for 1 hour (50% viability expected), and **F)** quiescent cells exposed to 68℃ for 1 hour (100% inviability expected). For the experiment, all cells were grown in rich media.

**Supplementary Figure 15.**
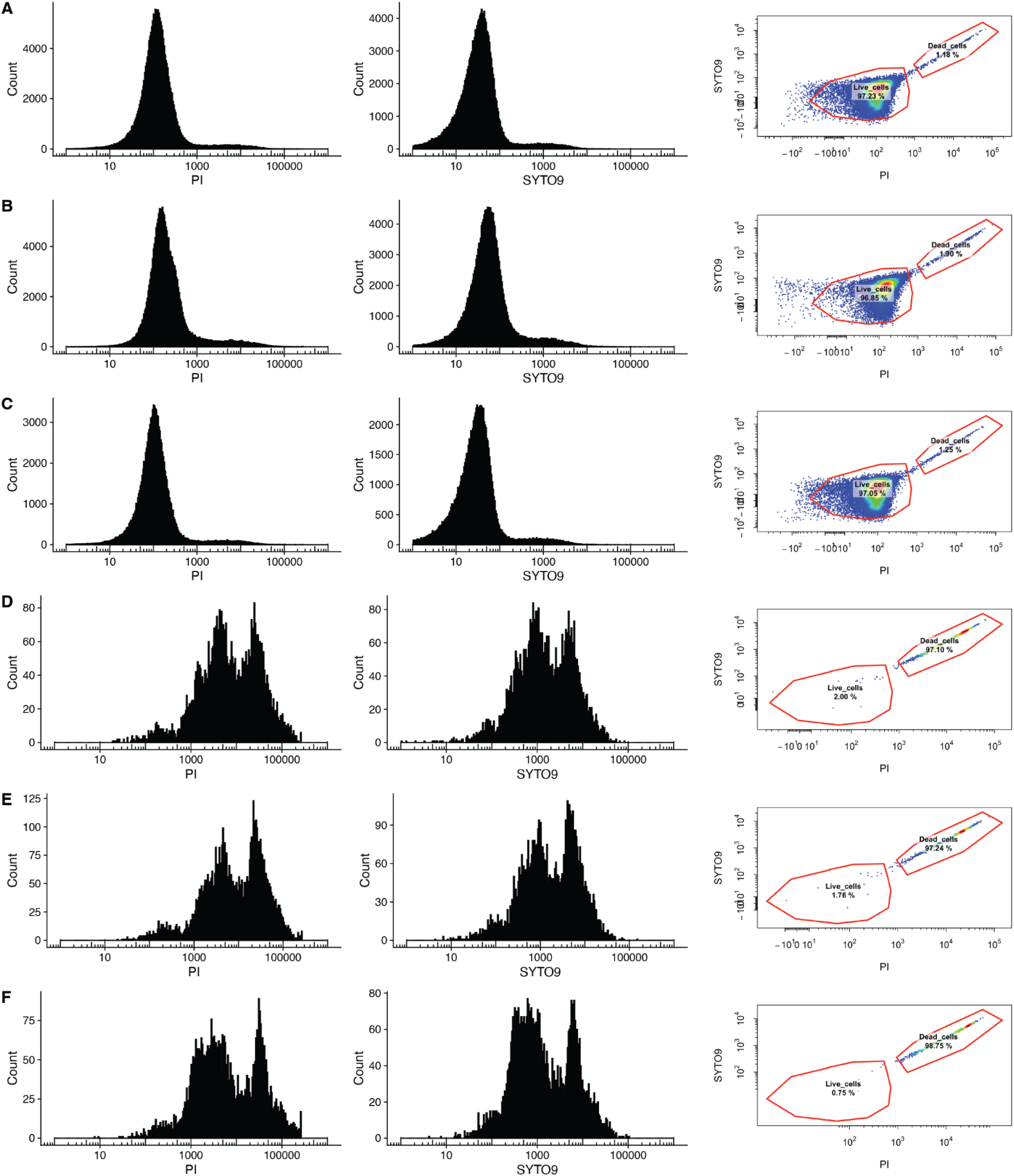
Cell viability quantification to validate accuracy of PI/SYTO9 staining quantification with caspofungin exposed proliferative cells. **A)** PI and SYTO9 distribution functions as well as the gated PI/SYTO9 2D quantification of untreated proliferative cells grown in rich media. **B)** PI and SYTO9 distribution functions as well as the gated PI/SYTO9 2D quantification of proliferative cells exposed to 0.001ug/mL caspofungin grown in rich media. **C)** PI and SYTO9 distribution functions as well as the gated PI/SYTO9 2D quantification of cells exposed to 0.01ug/mL caspofungin grown in rich media. **D)** PI and SYTO9 distribution functions as well as the gated PI/SYTO9 2D quantification of cells exposed to 0.1ug/mL caspofungin grown in rich media. **E)** PI and SYTO9 distribution functions as well as the gated PI/SYTO9 2D quantification of cells exposed to 1ug/mL caspofungin grown in rich media. **F)** PI and SYTO9 distribution functions as well as the gated PI/SYTO9 2D quantification of cells exposed to 10ug/mL caspofungin grown in rich media.

**Supplementary Figure 16.**
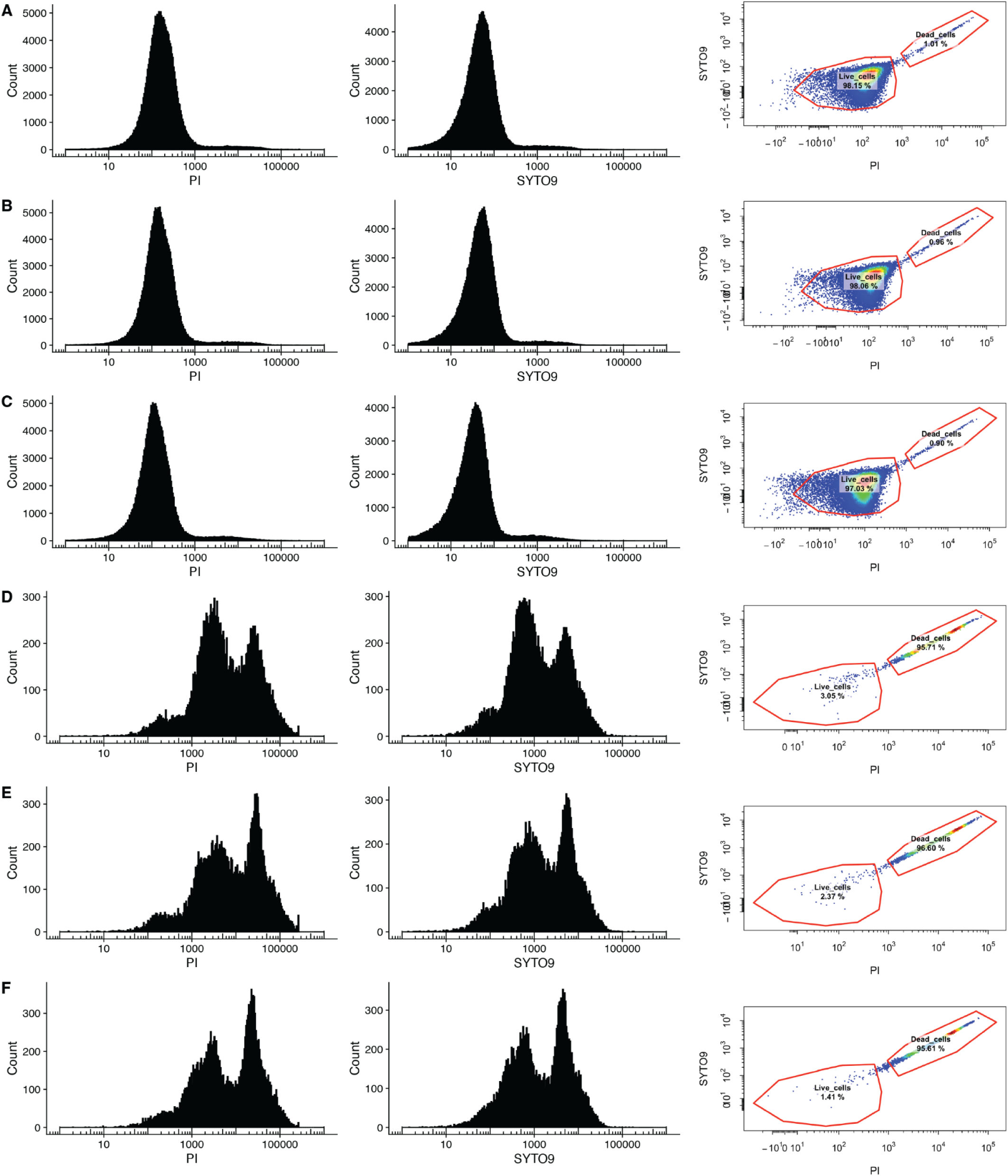
Cell viability quantification to validate accuracy of PI/SYTO9 staining quantification with micafungin exposed proliferative cells. **A)** PI and SYTO9 distribution functions as well as the gated PI/SYTO9 2D quantification of untreated proliferative cells grown in rich media. **B)** PI and SYTO9 distribution functions as well as the gated PI/SYTO9 2D quantification of proliferative cells exposed to 0.001ug/mL micafungin grown in rich media. **C)** PI and SYTO9 distribution functions as well as the gated PI/SYTO9 2D quantification of cells exposed to 0.01ug/mL micafungin grown in rich media. **D)** PI and SYTO9 distribution functions as well as the gated PI/SYTO9 2D quantification of cells exposed to 0.1ug/mL micafungin grown in rich media. **E)** PI and SYTO9 distribution functions as well as the gated PI/SYTO9 2D quantification of cells exposed to 1ug/mL micafungin grown in rich media. **F)** PI and SYTO9 distribution functions as well as the gated PI/SYTO9 2D quantification of cells exposed to 10ug/mL micafungin grown in rich media.

**Supplementary Figure 17.**
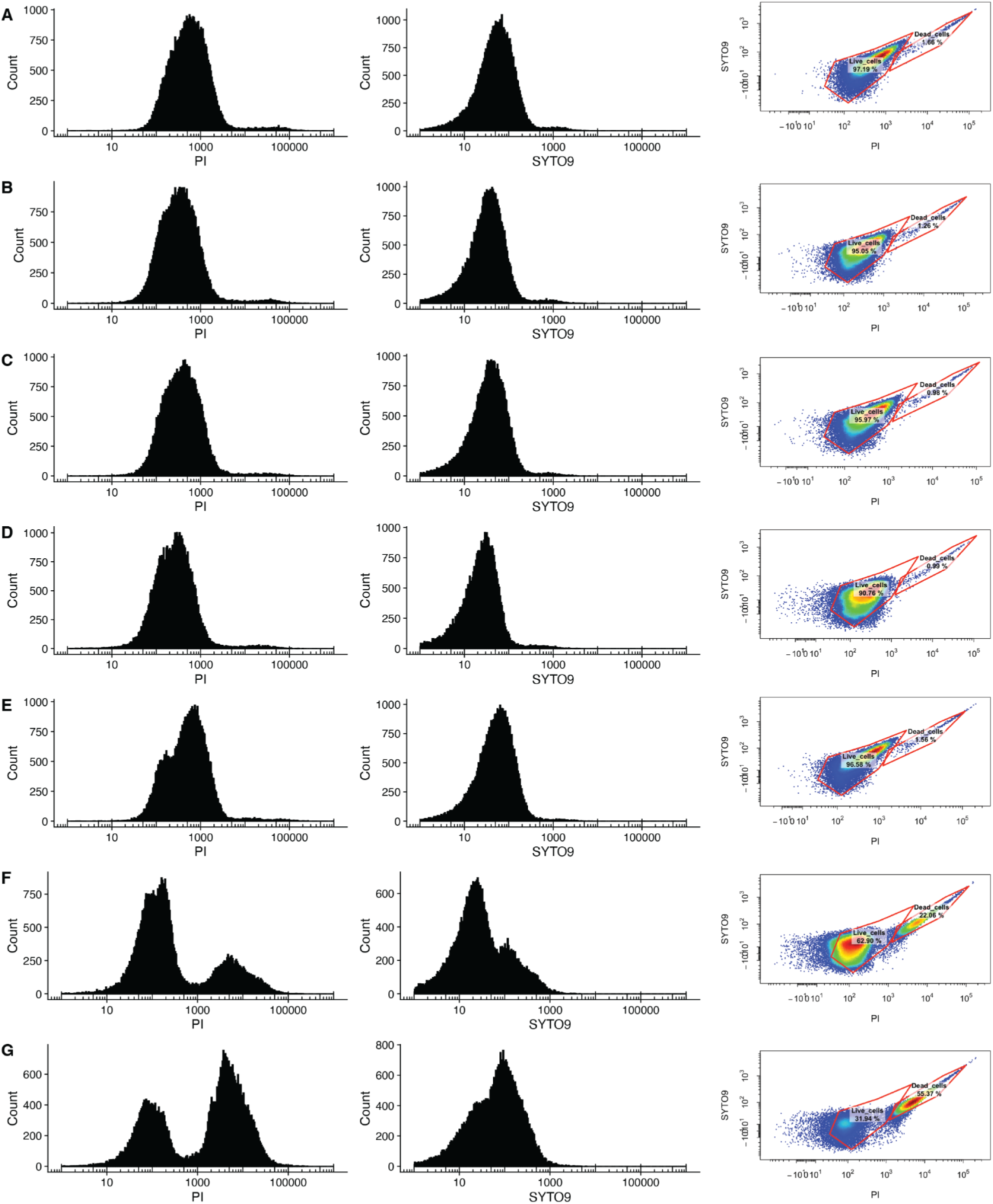
Cell viability quantification to validate accuracy of PI/SYTO9 staining quantification with amphotericin B exposed proliferative cells. **A)** PI and SYTO9 distribution functions as well as the gated PI/SYTO9 2D quantification of untreated proliferative cells grown in rich media. **B)** PI and SYTO9 distribution functions as well as the gated PI/SYTO9 2D quantification of proliferative cells exposed to 0.001ug/mL amphotericin B grown in rich media. **C)** PI and SYTO9 distribution functions as well as the gated PI/SYTO9 2D quantification of cells exposed to 0.01ug/mL amphotericin B grown in rich media. **D)** PI and SYTO9 distribution functions as well as the gated PI/SYTO9 2D quantification of cells exposed to 0.1ug/mL amphotericin B grown in rich media. **E)** PI and SYTO9 distribution functions as well as the gated PI/SYTO9 2D quantification of cells exposed to 1ug/mL amphotericin B grown in rich media. **F)** PI and SYTO9 distribution functions as well as the gated PI/SYTO9 2D quantification of cells exposed to 10ug/mL amphotericin B grown in rich media. **G)** PI and SYTO9 distribution functions as well as the gated PI/SYTO9 2D quantification of cells exposed to 20ug/mL amphotericin B grown in rich media.

**Supplementary Figure 18.**
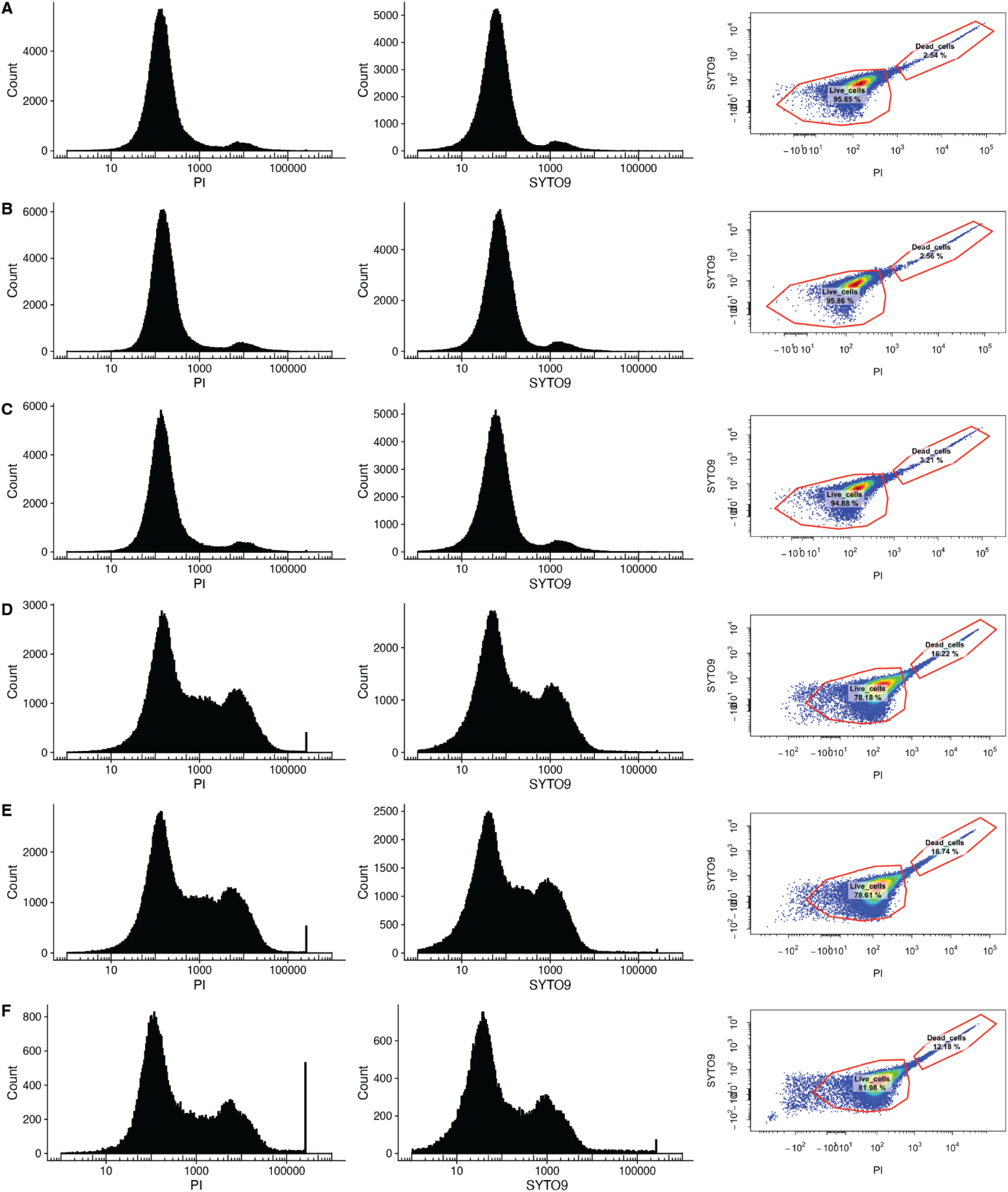
Cell viability quantification to validate accuracy of PI/SYTO9 staining quantification with caspofungin exposed quiescent cells. **A)** PI and SYTO9 distribution functions as well as the gated PI/SYTO9 2D quantification of untreated quiescent cells grown in rich media. **B)** PI and SYTO9 distribution functions as well as the gated PI/SYTO9 2D quantification of quiescent cells exposed to 0.001ug/mL caspofungin grown in rich media. **C)** PI and SYTO9 distribution functions as well as the gated PI/SYTO9 2D quantification of quiescent cells exposed to 0.01ug/mL caspofungin grown in rich media. **D)** PI and SYTO9 distribution functions as well as the gated PI/SYTO9 2D quantification of quiescent cells exposed to 0.1ug/mL caspofungin grown in rich media. **E)** PI and SYTO9 distribution functions as well as the gated PI/SYTO9 2D quantification of quiescent cells exposed to 1ug/mL caspofungin grown in rich media. **F)** PI and SYTO9 distribution functions as well as the gated PI/SYTO9 2D quantification of quiescent cells exposed to 10ug/mL caspofungin grown in rich media.

**Supplementary Figure 19.**
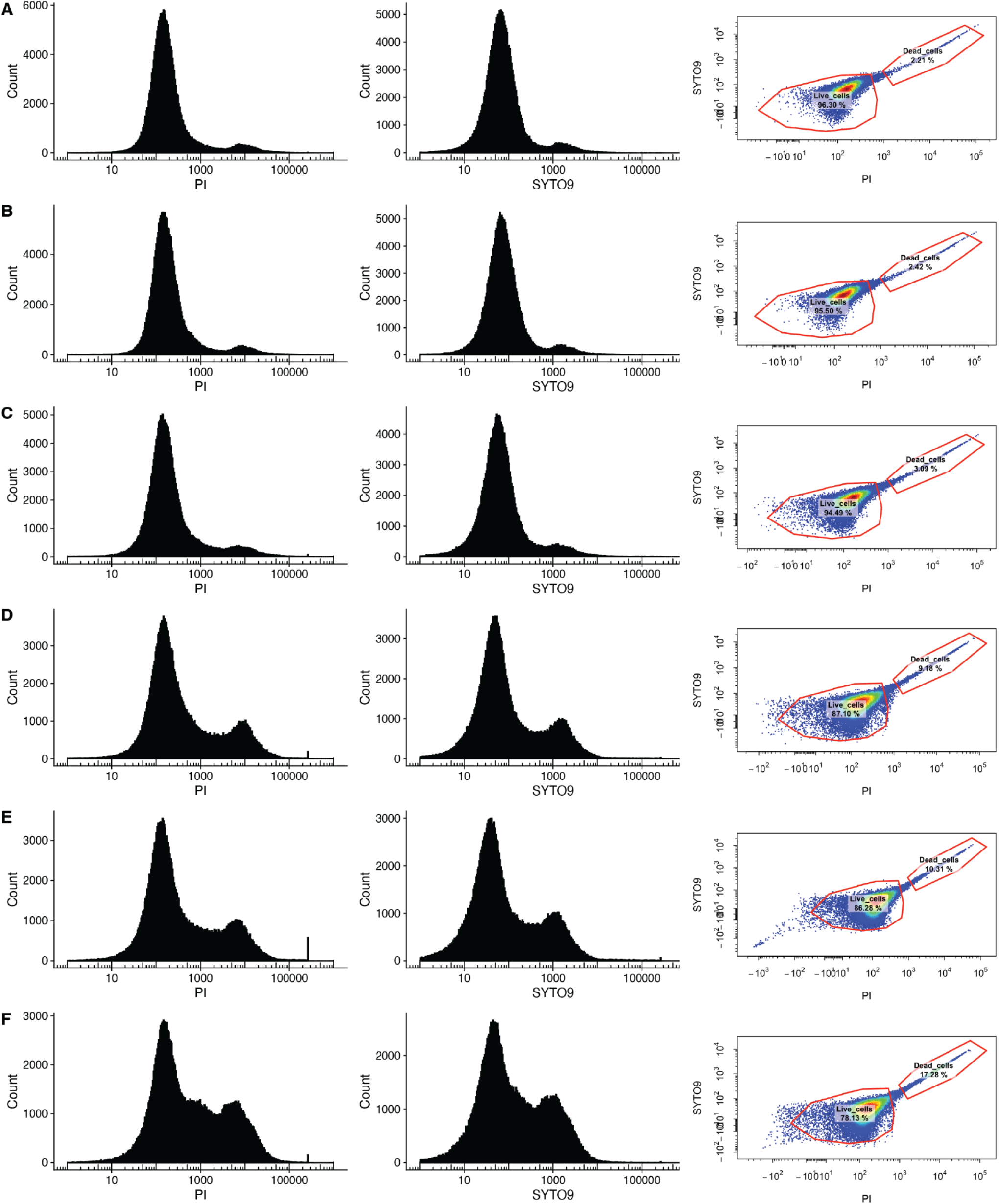
Cell viability quantification to validate accuracy of PI/SYTO9 staining quantification with micafungin exposed quiescent cells. **A)** PI and SYTO9 distribution functions as well as the gated PI/SYTO9 2D quantification of untreated quiescent cells grown in rich media. **B)** PI and SYTO9 distribution functions as well as the gated PI/SYTO9 2D quantification of quiescent cells exposed to 0.001ug/mL micafungin grown in rich media. **C)** PI and SYTO9 distribution functions as well as the gated PI/SYTO9 2D quantification of quiescent cells exposed to 0.01ug/mL micafungin grown in rich media. **D)** PI and SYTO9 distribution functions as well as the gated PI/SYTO9 2D quantification of quiescent cells exposed to 0.1ug/mL micafungin grown in rich media. **E)** PI and SYTO9 distribution functions as well as the gated PI/SYTO9 2D quantification of quiescent cells exposed to 1ug/mL micafungin grown in rich media. **F)** PI and SYTO9 distribution functions as well as the gated PI/SYTO9 2D quantification of quiescent cells exposed to 10ug/mL micafungin grown in rich media.

**Supplementary Figure 20.**
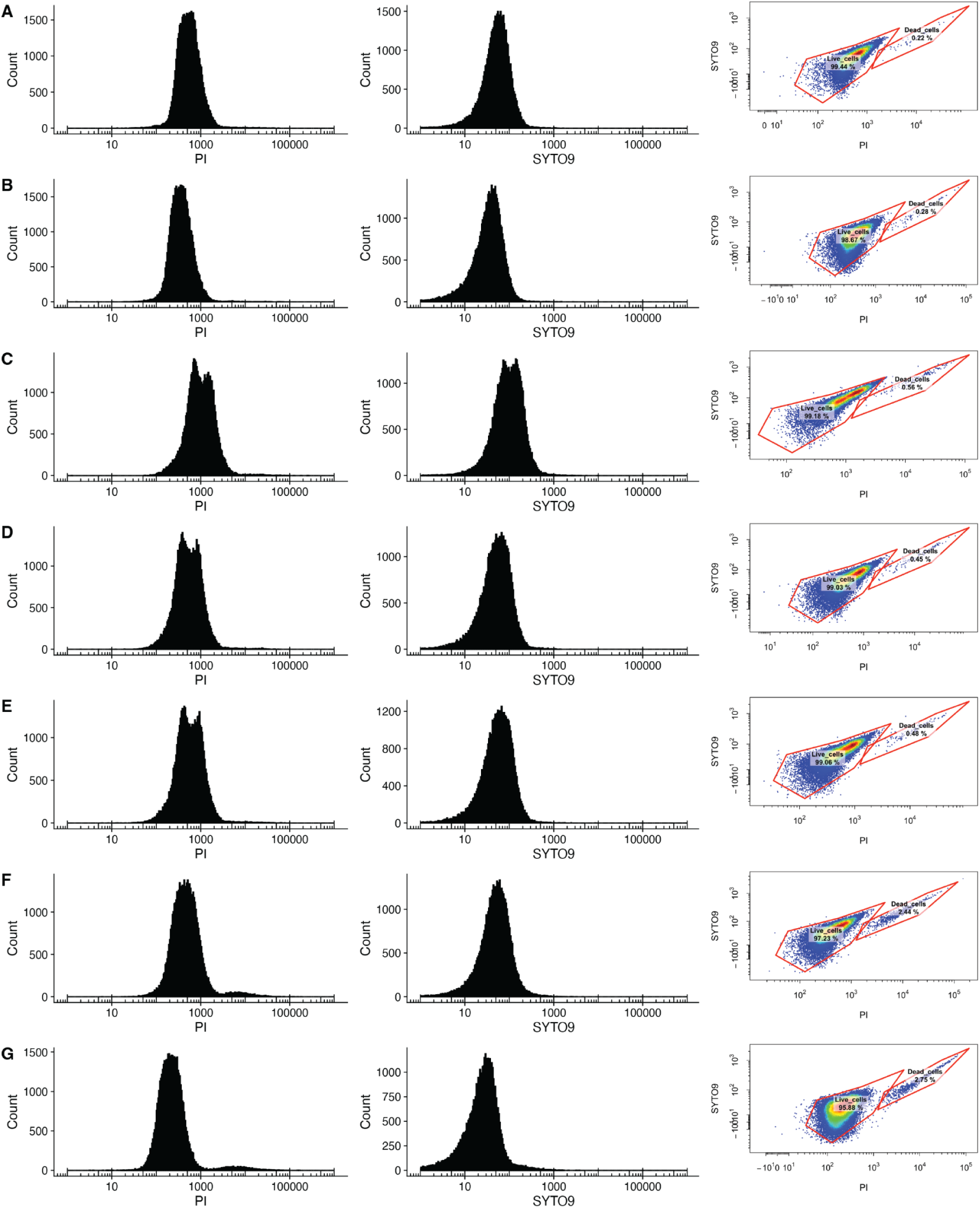
Cell viability quantification to validate accuracy of PI/SYTO9 staining quantification with amphotericin B exposed quiescent cells. **A)** PI and SYTO9 distribution functions as well as the gated PI/SYTO9 2D quantification of untreated quiescent cells grown in rich media. **B)** PI and SYTO9 distribution functions as well as the gated PI/SYTO9 2D quantification of quiescent cells exposed to 0.001ug/mL amphotericin B grown in rich media. **C)** PI and SYTO9 distribution functions as well as the gated PI/SYTO9 2D quantification of quiescent cells exposed to 0.01ug/mL amphotericin B grown in rich media. **D)** PI and SYTO9 distribution functions as well as the gated PI/SYTO9 2D quantification of quiescent cells exposed to 0.1ug/mL amphotericin B grown in rich media. **E)** PI and SYTO9 distribution functions as well as the gated PI/SYTO9 2D quantification of quiescent cells exposed to 1ug/mL amphotericin B grown in rich media. **F)** PI and SYTO9 distribution functions as well as the gated PI/SYTO9 2D quantification of quiescent cells exposed to 10ug/mL amphotericin B grown in rich media. PI and SYTO9 distribution functions as well as the gated PI/SYTO9 2D quantification of cells exposed to 20ug/mL amphotericin B grown in rich media.

## References

1. Bougnoux ME, Diogo D, François N, Sendid B, Veirmeire S, Colombel JF, et al. Multilocus sequence typing reveals intrafamilial transmission and microevolutions of Candida albicans isolates from the human digestive tract. J Clin Microbiol. 2006 May;44(5):1810–20.

2. Beigi RH, Meyn LA, Moore DM, Krohn MA, Hillier SL. Vaginal yeast colonization in nonpregnant women: a longitudinal study. Obstet Gynecol. 2004 Nov;104(5 Pt 1):926–30.

3. Jabra-Rizk MA, Kong EF, Tsui C, Nguyen MH, Clancy CJ, Fidel PL Jr, et al. Candida albicans pathogenesis: Fitting within the host-microbe damage response framework. Infect Immun. 2016 Oct 18;84(10):2724–39.

4. Katsipoulaki M, Stappers MHT, Malavia-Jones D, Brunke S, Hube B, Gow NAR. Candida albicans and Candida glabrata: global priority pathogens. Microbiol Mol Biol Rev. 2024 Jun 27;88(2):e0002123.

5. Tan CT, Xu X, Qiao Y, Wang Y. A peptidoglycan storm caused by β-lactam antibiotic’s action on host microbiota drives Candida albicans infection. Nat Commun. 2021 May 7;12(1):2560.

6. Cornely OA, Sprute R, Bassetti M, Chen SCA, Groll AH, Kurzai O, et al. Global guideline for the diagnosis and management of candidiasis: an initiative of the ECMM in cooperation with ISHAM and ASM. The Lancet Infectious Diseases. 2025 Feb 13;0(0).

7. Pfaller MA, Diekema DJ. Epidemiology of invasive candidiasis: a persistent public health problem. Clin Microbiol Rev. 2007 Jan;20(1):133–63.

8. WHO fungal priority pathogens list to guide research, development and public health action. World Health Organization; 2022.

9. Fisher MC, Alastruey-Izquierdo A, Berman J, Bicanic T, Bignell EM, Bowyer P, et al. Tackling the emerging threat of antifungal resistance to human health. Nat Rev Microbiol. 2022 Sep 29;20(9):557–71.

10. Rittershaus ESC, Baek SH, Sassetti CM. The normalcy of dormancy: common themes in microbial quiescence. Cell Host Microbe. 2013 Jun 12;13(6):643–51.

11. Sun S, Gresham D. Cellular quiescence in budding yeast. Yeast. 2021 Jan;38(1):12–29.

12. Trutneva KA, Shleeva MO, Demina GR, Vostroknutova GN, Kaprelyans AS. One-Year Old Dormant, “Non-culturable” Mycobacterium tuberculosis Preserves Significantly Diverse Protein Profile. Front Cell Infect Microbiol. 2020 Jan 31;10:26.

13. Parrish NM, Dick JD, Bishai WR. Mechanisms of latency in Mycobacterium tuberculosis. Trends Microbiol. 1998 Mar 1;6(3):107–12.

14. Laporte D, Gouleme L, Jimenez L, Khemiri I, Sagot I. Mitochondria reorganization upon proliferation arrest predicts individual yeast cell fate. Elife. 2018 Oct 9;7:e35685.

15. Sagot I, Laporte D. The cell biology of quiescent yeast - a diversity of individual scenarios. J Cell Sci. 2019 Jan 2;132(1).

16. Jacquel B, Aspert T, Laporte D, Sagot I, Charvin G. Monitoring single-cell dynamics of entry into quiescence during an unperturbed life cycle. Elife. 2021 Nov 1;10.

17. Enriquez-Hesles E, Smith DL Jr, Maqani N, Wierman MB, Sutcliffe MD, Fine RD, et al. A cell-nonautonomous mechanism of yeast chronological aging regulated by caloric restriction and one-carbon metabolism. J Biol Chem. 2021 Jan;296(100125):100125.

18. Klosinska MM, Crutchfield CA, Bradley PH, Rabinowitz JD, Broach JR. Yeast cells can access distinct quiescent states. Genes Dev. 2011;25(4):336–49.

19. De Virgilio C. The essence of yeast quiescence. FEMS Microbiol Rev. 2012;36(2):306–39.

20. Gangloff S, Arcangioli B. DNA repair and mutations during quiescence in yeast. FEMS Yeast Res. 2017 Jan 1;17(1).

21. Jarvis B, Figgitt DP, Scott LJ. micafungin. Drugs. 2004;64(9):969–82; discussion 983–4.

22. Maji A, Soutar CP, Zhang J, Lewandowska A, Uno BE, Yan S, et al. Tuning sterol extraction kinetics yields a renal-sparing polyene antifungal. Nature. 2023 Nov;623(7989):1079–85.

23. Chandrasekar PH, Sobel JD. micafungin: a new echinocandin. Clin Infect Dis. 2006 Apr 15;42(8):1171–8.

24. Wei W, Nurse P, Broek D. Yeast cells can enter a quiescent state through G1, S, G2, or M phase of the cell cycle. Cancer Res. 1993 Apr 15;53(8):1867–70.

25. Laporte D, Lebaudy A, Sahin A, Pinson B, Ceschin J, Daignan-Fornier B, et al. Metabolic status rather than cell cycle signals control quiescence entry and exit. J Cell Biol. 2011 Mar 21;192(6):949–57.

26. Delarue M, Brittingham GP, Pfeffer S, Surovtsev IV, Pinglay S, Kennedy KJ, et al. mTORC1 Controls Phase Separation and the Biophysical Properties of the Cytoplasm by Tuning Crowding. Cell. 2018 Jul 12;174(2):338–49.e20.

27. Munder MC, Midtvedt D, Franzmann T, Nüske E, Otto O, Herbig M, et al. A pH-driven transition of the cytoplasm from a fluid- to a solid-like state promotes entry into dormancy. Vol. 5, eLife. 2016.

28. Levy SF, Ziv N, Siegal ML. Bet hedging in yeast by heterogeneous, age-correlated expression of a stress protectant. PLoS Biol. 2012 May 8;10(5):e1001325.

29. Witkowski B, Lelièvre J, Barragán MJL, Laurent V, Su XZ, Berry A, et al. Increased tolerance to artemisinin in Plasmodium falciparum is mediated by a quiescence mechanism. Antimicrob Agents Chemother. 2010 May;54(5):1872–7.

30. Tang H, Liang J, He BZ. An optimized LIVE/DEAD assay using flow cytometry to quantify post-stress and antifungal-treatment survival in diverse yeasts. bioRxiv. p. 2025.04.14.648826.

31. Li P, Seneviratne CJ, Luan Q, Jin L. Proteomic Analysis of caspofungin-Induced Responses in Planktonic Cells and Biofilms of Candida albicans. Front Microbiol. 2021;12:639123.

32. Shi L, Sutter BM, Ye X, Tu BP. Trehalose is a key determinant of the quiescent metabolic state that fuels cell cycle progression upon return to growth. Mol Biol Cell. 2010 Jun 15;21(12):1982–90.

33. Persson LB, Ambati VS, Brandman O. Cellular Control of Viscosity Counters Changes in Temperature and Energy Availability. Cell. 2020 Dec 10;183(6):1572–85.e16.

34. Monterroso B, Margolin W, Boersma AJ, Rivas G, Poolman B, Zorrilla S. Macromolecular crowding, phase separation, and homeostasis in the orchestration of bacterial cellular functions. Chem Rev. 2024 Feb 28;124(4):1899–949.

35. LaFleur MD, Kumamoto CA, Lewis K. Candida albicans biofilms produce antifungal-tolerant persister cells. Antimicrob Agents Chemother. 2006 Nov;50(11):3839–46.

36. Gray JV, Petsko GA, Johnston GC, Ringe D, Singer RA, Werner-Washburne M. “Sleeping beauty”: quiescence in Saccharomyces cerevisiae. Microbiol Mol Biol Rev. 2004 Jun;68(2):187–206.

37. Kussell E, Leibler S. Phenotypic diversity, population growth, and information in fluctuating environments. Science. 2005 Sep 23;309(5743):2075–8.

38. Balaban NQ, Merrin J, Chait R, Kowalik L, Leibler S. Bacterial persistence as a phenotypic switch. Science. 2004 Sep 10;305(5690):1622–5.

39. Kramara J, Kim MJ, Ollinger TL, Ristow LC, Wakade RS, Zarnowski R, et al. Systematic analysis of the Candida albicans kinome reveals environmentally contingent protein kinase-mediated regulation of filamentation and biofilm formation in vitro and in vivo. MBio. 2024 Aug 14;15(8):e0124924.

40. Galanti L, Shasha D, Gunsalus KC. Pheniqs 2.0: accurate, high-performance Bayesian decoding and confidence estimation for combinatorial barcode indexing. BMC Bioinformatics. 2021 Jul 2;22(1):359.

41. GENEFLOW: Genomic Examination and Nucleotide Evaluation For Laboratory Operations Workflow. Github: https://github.com/gencorefacility/GENEFLOW

42. Keegan S, Fenyö D, Holt LJ. GEMspa: a Napari plugin for analysis of single particle tracking data. bioRxiv. 2023. p. 2023.06.26.546612.

43. Hirakawa MP, Martinez DA, Sakthikumar S, Anderson MZ, Berlin A, Gujja S, et al. Genetic and phenotypic intra-species variation in Candida albicans. Genome Res. 2015 Mar;25(3):413–25.

